# Effects of Polyphosphate metabolism Mutations on Biofilm, Capsule Formation and Virulence Traits in Hypervirulent ST23 *Klebsiella pneumoniae* SGH10

**DOI:** 10.1101/2023.09.15.557991

**Authors:** Diego Rojas, Andrés E. Marcoleta, Matías Gálvez, Macarena A. Varas, Mauricio Díaz, Mauricio Hernández, Cristian Vargas, Guillermo Nourdin, Elard Koch, Pablo Saldivia, Jorge Vielma, Yunn-Hwen Gan, Yahua Chen, Nicolás Guiliani, Francisco P. Chávez

## Abstract

The emergence of hypervirulent *Klebsiella pneumoniae* (hvKP) strains poses a significant threat to public health due to their high mortality rates and propensity to cause severe community-acquired infections in otherwise healthy individuals. The ability of hvKP to form biofilms and produce a protective capsule contributes to its enhanced virulence and is a significant challenge to effective antibiotic treatment. Therefore, understanding the molecular mechanisms underlying hvKP virulence and biofilm formation is crucial for developing new therapeutic strategies. Polyphosphate Kinase 1 (PPK1) is an enzyme responsible for inorganic polyphosphate synthesis and plays a vital role in regulating various physiological processes in bacteria. In this study, we investigated the impact of polyP metabolism on the biofilm and capsule formation and virulence traits in hvKP using *Dictyostelium discoideum* amoeba as a model host. We found that the PPK1 null-mutant was impaired in biofilm and capsule formation and showed attenuated virulence in *D. discoideum* compared to the wild-type strain. We performed a shotgun proteomic analysis of the PPK1 mutant and wild-type strain to gain further insight into the underlying molecular mechanism. The results revealed that the PPK1 mutant had a differential expression of proteins (DEP) involved in capsule synthesis (Wzi - Ugd), biofilm formation (MrkC-D-H), synthesis of the colibactin genotoxin precursor (ClbB), as well as proteins associated with the synthesis and modification of lipid A (ArnB -LpxC - PagP). These proteomic findings corroborate the phenotypic observations and indicate that the PPK1 mutation is associated with impaired biofilm and capsule formation and attenuated virulence in hypervirulent *K. pneumoniae*. Overall, our study highlights the importance of polyP synthesis in regulating extracellular biomolecules and virulence in *K. pneumoniae* and provides insights into potential therapeutic targets for treating *K. pneumoniae* infections.

## INTRODUCTION

Hypervirulent *Klebsiella pneumoniae* (hvKP) strains have emerged as a significant public health threat due to their ability to cause severe and often fatal infections in healthy individuals ^1^. Unlike traditional *K. pneumoniae* strains primarily cause hospital-acquired infections in immunocompromised patients, hvKP strains can cause community-acquired invasive infections in healthy individuals without any underlying medical conditions ^2^. These infections can lead to various clinical manifestations, such as pyogenic liver abscesses (PLA), meningitis, endophthalmitis, necrotizing fasciitis, and sepsis, with mortality rates as high as 50% ^3^. Moreover, the dynamic nature of the *K. pneumoniae* genome and the acquisition of virulence and resistance factors through horizontal gene transfer have facilitated the rapid evolution of new *K. pneumoniae* strains exhibiting both hypervirulence (hv) and multidrug resistance (MDR), rendering them resistant to all conventional treatment methods ^4^. Hallmark traits identified in hvKP strains include the production of a thick and prominent capsule and a hypermucoviscous phenotype ^2^. These traits are associated with a virulence plasmid (∼200 kb), which harbors two genes encoding transcriptional regulators (*rmpA1* and *rmpA2*) that activate the expression of the 20-gene cps operon/locus. Additionally, the *rmpD* gene encodes a small protein required for hypermucoviscosity (HMV) ^5^, promoting increased capsular polysaccharide chain length ^6^. Furthermore, the *iucA* gene, which codes for the aerobactin siderophore, is also involved in hvKP virulence ^7^.

Surface structures, such as lipopolysaccharide (LPS) and fimbriae, frequently encoded in the core genome, play a crucial role in the invasion of epithelial cells and biofilm formation on biotic and abiotic surfaces. Notably, the expression of type I and III fimbriae has been associated with the virulence of hvKP ^8–11^. Genetic experiments conducted in various models have revealed that biofilm and capsule formation, particularly with specific capsular serotypes like K1 and K2, are the main mechanisms employed by hvKP to evade the immune response ^12,13^. Therefore, targeting biofilm and capsule production could be a promising strategy to control *K. pneumoniae* infections. However, further studies are necessary to elucidate the molecular mechanisms regulating biofilm formation and capsule production in this hvKP.

Understanding these mechanisms would be valuable in the search for treatments against Sequence Type 23 (ST23), which encompasses most of the hvKP strains associated with severe infections in humans, as well as emerging lineages exhibiting the convergence of hypervirulence and resistance to carbapenems and other last-line antibiotics ^14^. Notably, strain SGH10 has been described as the best representative of hvKP ST23 strains and is a suitable model for studying their pathogenesis ^15^. Hence, exploring novel approaches and targets to combat these hypervirulent strains is crucial. Inorganic polyphosphate (polyP) is ubiquitous in all living organisms, from bacteria to humans, and plays a critical role in various cellular processes.

In bacteria, polyP metabolism has been shown to contribute to stress response, antibiotic resistance, persistence, and virulence ^16–18^ For example, in *Pseudomonas aeruginosa*, polyP is involved in biofilm formation and quorum sensing, contributing to its pathogenicity ^19–21^. Similarly, *Salmonella enterica* utilizes polyP for intracellular survival ^22,23^, and recent studies have highlighted the significance of polyP and the enzyme PPK1 in intracellular survival within macrophages and modulation of the immune response ^24,25^. In the context of *K. pneumoniae* and polyP metabolism, the effect of gallein, an inhibitor of polyP synthesis, has been studied, revealing a concentration-dependent decrease in biofilm formation ^26^. hvKP strains are associated with enhanced virulence and have been observed to form biofilms and produce a protective capsule, contributing to their pathogenicity ^27^. However, the precise mechanism by which polyP metabolism and PPK1 influence *Klebsiella* virulence remains poorly understood. Here, we investigated the impact of PPK1 mutation on hvKP biofilm, capsule formation, and virulence traits using *Dictyostelium discoideum* amoeba as a model host.

## RESULTS

### Deleting polyP metabolism genes does not affect viability and differentially alters polyP accumulation

To investigate polyP metabolism and its influence on virulence traits, gene knockout mutants were generated in both the synthesis (τι*ppk1*) and degradation (τι*ppx*) of polyP, as well as the double mutant (τι*ppk1*-τι*ppx*) in the hypervirulent strain ST23 K. pneumoniae SGH10 (Fig. 1S). Upon acquisition and genomic sequence confirmation via Illumina, the obtained mutants were analyzed for their ability to produce capsules and biofilms and in virulence assays using the social amoeba *D. discoideum*. We first characterized the growth of polyP metabolism mutants (Fig. S1) under optimal conditions (LB broth, 37°C, 180 rpm) and minimal medium (MOPS). Deleting *ppk1, ppx*, and *ppk1*-*ppx* genes did not affect growth under normal conditions (Fig. S2A). However, under nutrient-deprived conditions, a delay of approximately 2 h in reaching exponential growth was observed in the τι*ppk1* and τι*ppk1*-τι*ppx* mutants (Fig. S2B). This delay was not observed in the τι*ppx* mutant, suggesting that the absence of PPK1 specifically affects normal growth under nutrient-deficient conditions. Furthermore, adding different concentrations of polyP 150-mer can restore growth under minimal nutrient conditions (Fig. S2B).

PPK1 is the main enzyme responsible for synthesizing polyP in bacterial models such as *P. aeruginosa* and *Acinetobacter baumannii* ^23,28^. However, its role in *K. pneumoniae* has not been extensively studied. Therefore, polyP levels were evaluated under favorable conditions for its accumulation. As expected, we found that polyP levels significantly decreased in the Δ*ppk1* and Δ*ppk1*-Δ*ppx* mutants compared to the wild-type (WT) strain (Fig. 1A). No significant change in polyP levels was observed in the Δ*ppx* mutant compared to the wild type. Furthermore, polyP accumulation was visualized using confocal microscopy with DAPI staining ^29^(Fig. 1B). Bacteria were cultured in a rich medium and transferred to a nutrient-poor medium (MOPS) for two hours. Compared to the Δ*ppx* and WT strains, a significantly lower number of bacteria in the Δ*ppk1* and Δ*ppk1*-Δ*ppx* mutants was observed, which correlated with the growth curve delay observed in the minimal medium (Fig. S1B). Additionally, we noted a decrease in the formation and accumulation of polyP granules, as evidenced by the reduction in the fluorescence intensity of DAPI-polyP in Δ*ppk1* and Δ*ppk1*-Δ*ppx* strains compared to the WT and Δ*ppx* strains. Mesalamine, a PPK1 inhibitor ^30^, also reported a similar effect. Our results suggest that PPK1 is the primary enzyme responsible for polyP synthesis in *K. pneumoniae*, and the absence of exopolyphosphatase (PPX) does not significantly increase polyP synthesis or accumulation.

**Figure 1.**
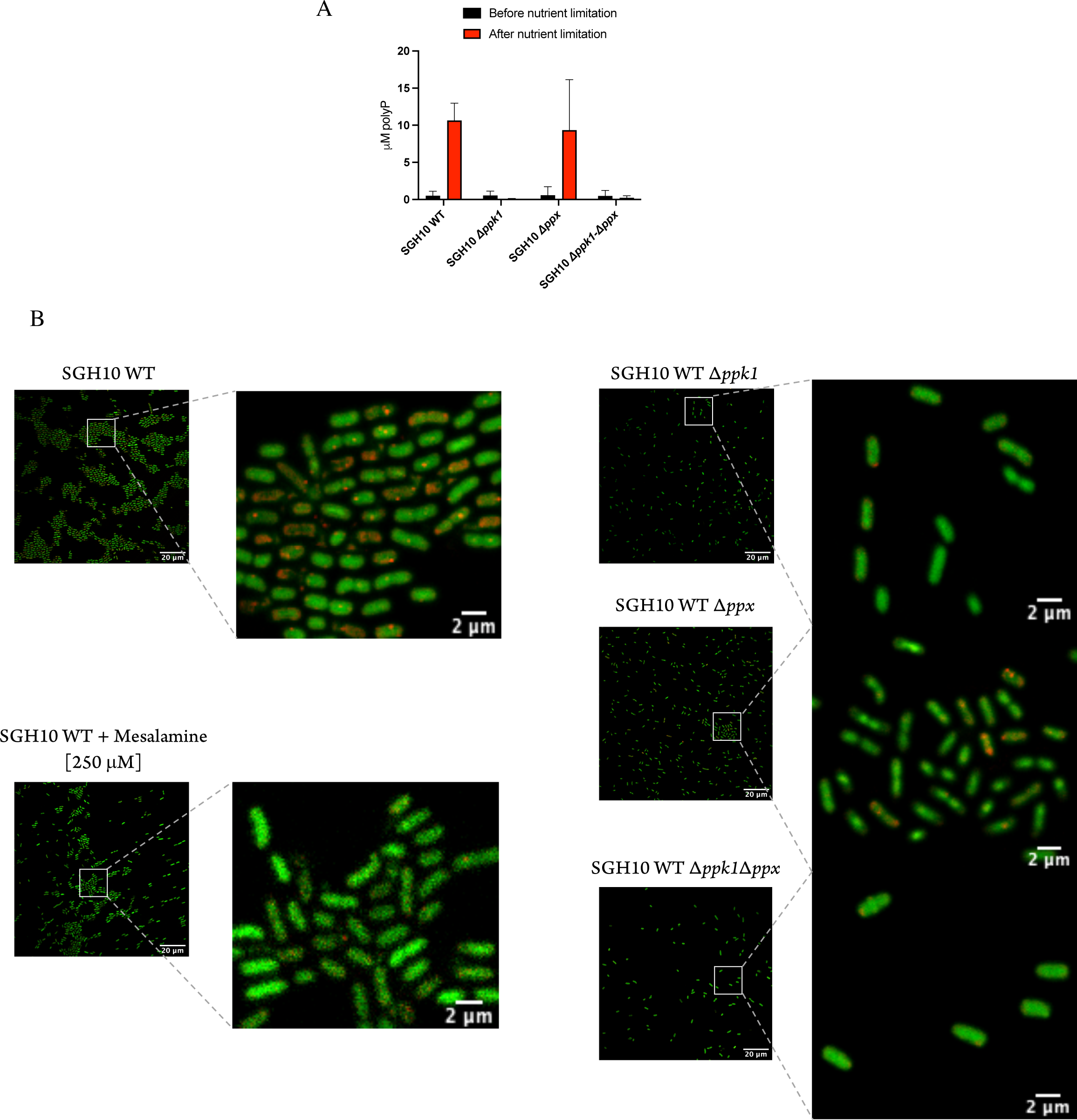
PolyP accumulation under nutrient deprivation. (A) An overnight culture was diluted 1:100 in 10 mL of LB broth and allowed to grow with agitation until the mid-logarithmic phase (approximately 2 h). 1 mL of culture cells were pelleted and washed three times with MOPS medium (0.4% glucose and 0.1 mM K2HPO4), then incubated with agitation for 2 h at 37°C. The level of polyP was quantified before and after the nutritional shift. (B) The culture was set in the dark with DAPI (5 μg/mL) for 30 min and then mounted for observation by confocal microscopy using a 63X oil immersion objective. DNA was visualized using an excitation wavelength of 378 nm and an emission wavelength of 456 nm. PolyP was detected with excitation at 415 nm and emission at 550 nm. DAPI-DNA is represented in green, while DAPI-polyP is indicated in red for clarity in our visual representation.

### PolyP metabolism mutants show altered capsule and mucoviscosity

The capsule is a critical virulence factor in hypervirulent strains, and this pathotype has been associated with a small group of capsular serotypes, namely K1 and K2, which are relevant for the pathogenicity of a wide range of strains ^31,32^. Also, hvKP frequently exhibits an HMV phenotype, which has recently been linked to the polysaccharide length rather than its modification ^6^. HMV strains show poor sedimentation during low-speed centrifugation, resulting in a turbid supernatant. This unique characteristic allows turbidity measurements after centrifugation as a quantitative indicator of HMV ^5^. Here, mucoviscosity was quantified and a significant reduction in all mutants involved in polyP metabolism was observed when compared to the WT strain (Fig. 2A). Furthermore, we assessed the role of uronic acids (UA) as essential components found in numerous capsules, traditionally serving as an indicator of capsule content ^33^. Interestingly, results showed a slight but significant reduction in capsule production in the Δ*ppk1* and Δ*ppk1*-Δ*ppx* (Fig. 2B). These findings highlight the importance of PPK1 enzyme activity and polyP levels in regulating capsule formation, further underscoring their role in the virulence of hypervirulent strains. To elucidate whether the observed disparities in mucoviscosity and capsule quantity among the studied mutant strains are attributable to any surface modifications, we conducted a comprehensive assessment of these strains via scanning electron microscopy (SEM). Previous studies have indicated that the capsule of the SGH10 strain extends from the surface in multiple uniformly distributed directions ^34–36^. We observed variations in the amount of capsular polysaccharide layer in the Δ*ppk1* and Δ*ppk1*-Δ*ppx* mutants, which is evident in the WT strain (Fig. 2C).

**Figure 2.**
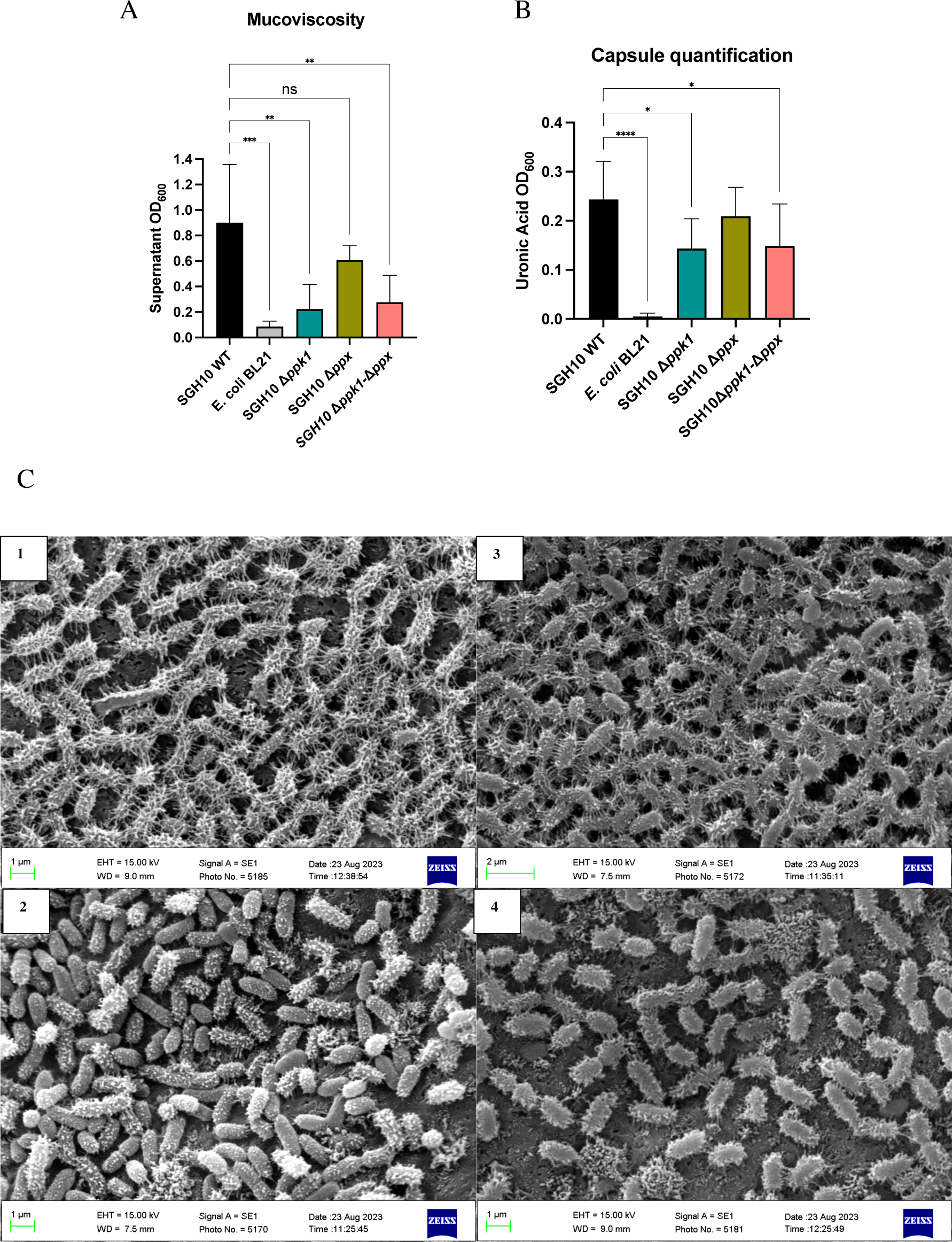
Capsule production and mucoviscosity in polyP-deficient *K. pneumoniae* mutants. (A) Strain mucoid was measured by low-speed centrifugation (3000 rpm for 10 min) in LB. The optical density at 600 nm (OD600) of the supernatant above the pellet was measured and mean ± SD was plotted (n = 3). (B) Capsule production of polyP metabolism mutants (Δ*ppk1*, Δ ppx and Δ*ppk1-*Δ*ppx*). Uronic acid quantification analysis as described in Materials and Methods. One-way ANOVA tests were performed for statistical analyses comparing the WT strain to the indicated mutants. *, P < 0.05. ****, P <0.0001. (C) Scanning electron microscopy images of cells fixed in glutaraldehyde 2.5% and 0.15% ruthenium red, obtained from log-phase cultures of SGH10 WT (1), SGH10 Δ*ppk1* (2), SGH10 Δ*ppx* (3) and SGH10 Δ*ppk1*-Δ*ppx* (4)cells. The scale bar is 1 μm.

### PPK1 plays a crucial role in biofilm formation

Biofilm formation is a critical aspect of bacterial virulence and persistence in the host, facilitated by fimbrial adhesins ^37,38^. These structures have significant clinical implications as they contribute to essential processes such as antimicrobial resistance and chronic endurance of infectious diseases. Recently, it has been reported that mesalamine, a drug used to treat ulcerative colitis, selectively inhibits PPK1 in gastrointestinal tract bacteria, decreasing biofilm formation and virulence ^30^. To investigate whether mutants involved in polyP metabolism exhibit alterations in biofilm formation, we conducted a biomass quantification assay using crystal violet staining (0.1%). The Δ*ppk1* and Δ*ppk1*-Δ*ppx* mutants showed a 66% and 56% decrease in biofilm formation compared to the WT strain (Fig. 3A). This decrease was not observed in the Δ*ppx* mutant, suggesting that the absence of PPK1 directly affects biofilm formation. Furthermore, we used mesalamine to validate this observation and observed a 76% reduction in biofilm biomass compared to the WT strain (Fig. 3A). Another approach to quantify biofilm formation was performed by measuring extracellular polymeric substances (EPS) levels such as curli and cellulose. For this purpose, we employed the Ebbabiolight 680 probe ^39^ an optotracer that binds to curli and cellulose, increasing its fluorescence (Relative fluorescence units). A consistent decrease in biofilm formation in Δ*ppk1* and Δ*ppk1*-Δ*ppx* was observed (Fig. 3B), as quantified by crystal violet staining. The observation of a 24-hour biofilm grown on a tilted 96-well microtiter plate under the confocal microscope further confirmed that the Δ*ppk1* and Δ*ppk1*-Δ*ppx* mutants exhibited reduced biofilm biomass compared to the WT and Δ*ppx* (Fig 3C). To better understand the relationship between polyP metabolism and biofilm formation, we conducted a macrobiofilm formation assay in a salt-free LB medium supplemented with Thioflavin S, which selectively binds to amyloid fibers. In the SGH10 WT strain, we observed a macrobiofilm with a central mucoid region, dense under light, displaying regular concentric rings. In the Δ*ppk1* mutant, we observed a reduction in mucoidness in the center of the colony, the absence of concentric rings, and a smoother appearance with irregular edges. No significant differences were observed in the Δ*ppx* mutant compared to the wild-type strain, as it exhibited a central mucoid ring and regular concentric rings with a smooth edge. In Δp*pk1*-Δ*ppx*, we observed a reduction in the mucoid ring in the center of the macrobiofilm, along with the absence of regular concentric rings. However, the edges were like the WT strain, presenting an intermediate phenotype between the wild-type strain and the Δ*ppk1* mutant. In conclusion, our findings demonstrate that PPK1 plays a crucial role in biofilm formation, as evidenced by the significant reduction in biofilm biomass observed in the Δ*ppk1* and Δ*ppk1*-Δ*ppx* mutants. The inhibitory effect of mesalamine on biofilm formation further supports the involvement of PPK1 in this process. These results provide valuable insights into the regulatory mechanisms of biofilm formation and highlight potential therapeutic targets for combating biofilm-associated infections ^37,38,30^.

**Figure 3.**
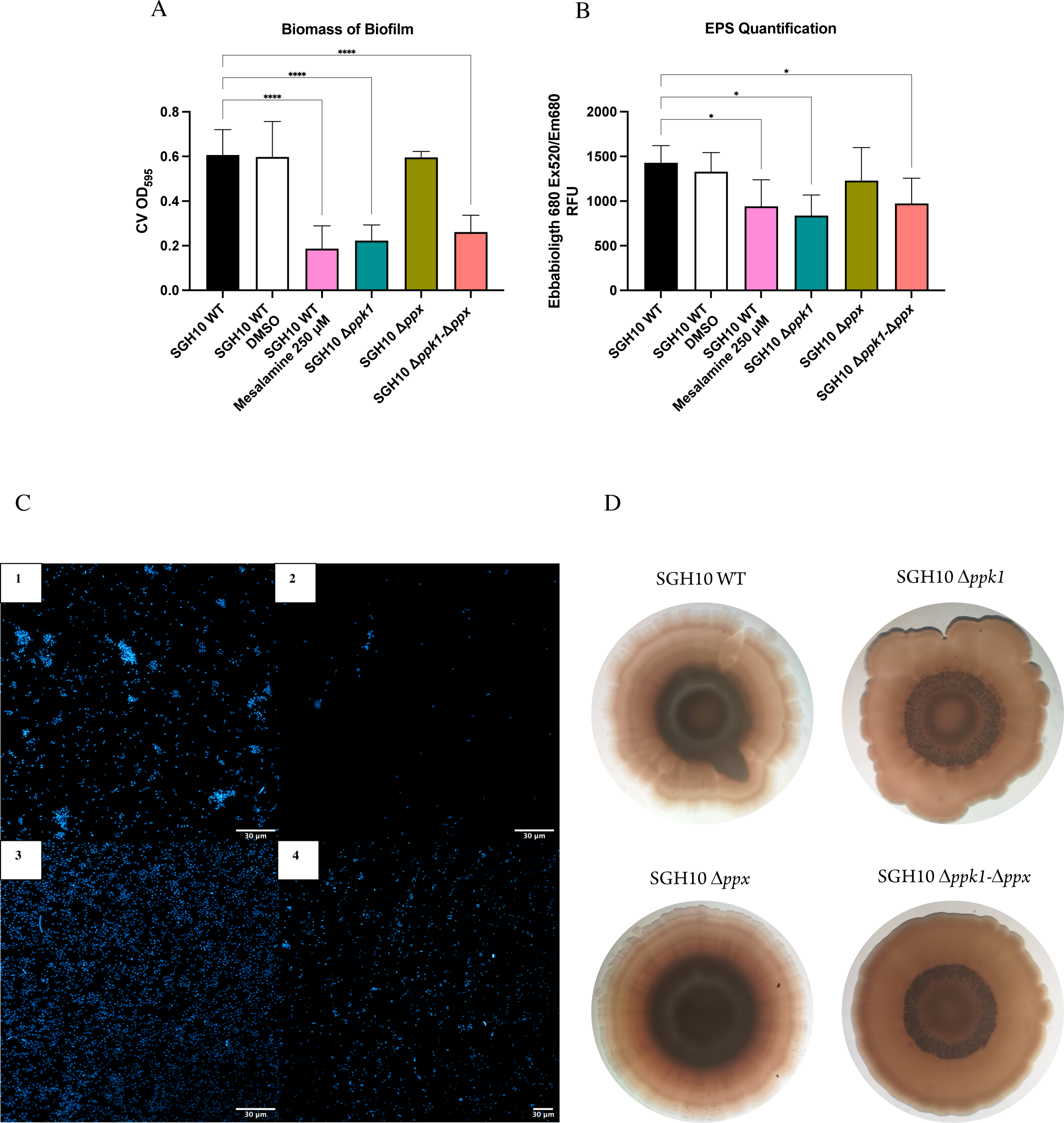
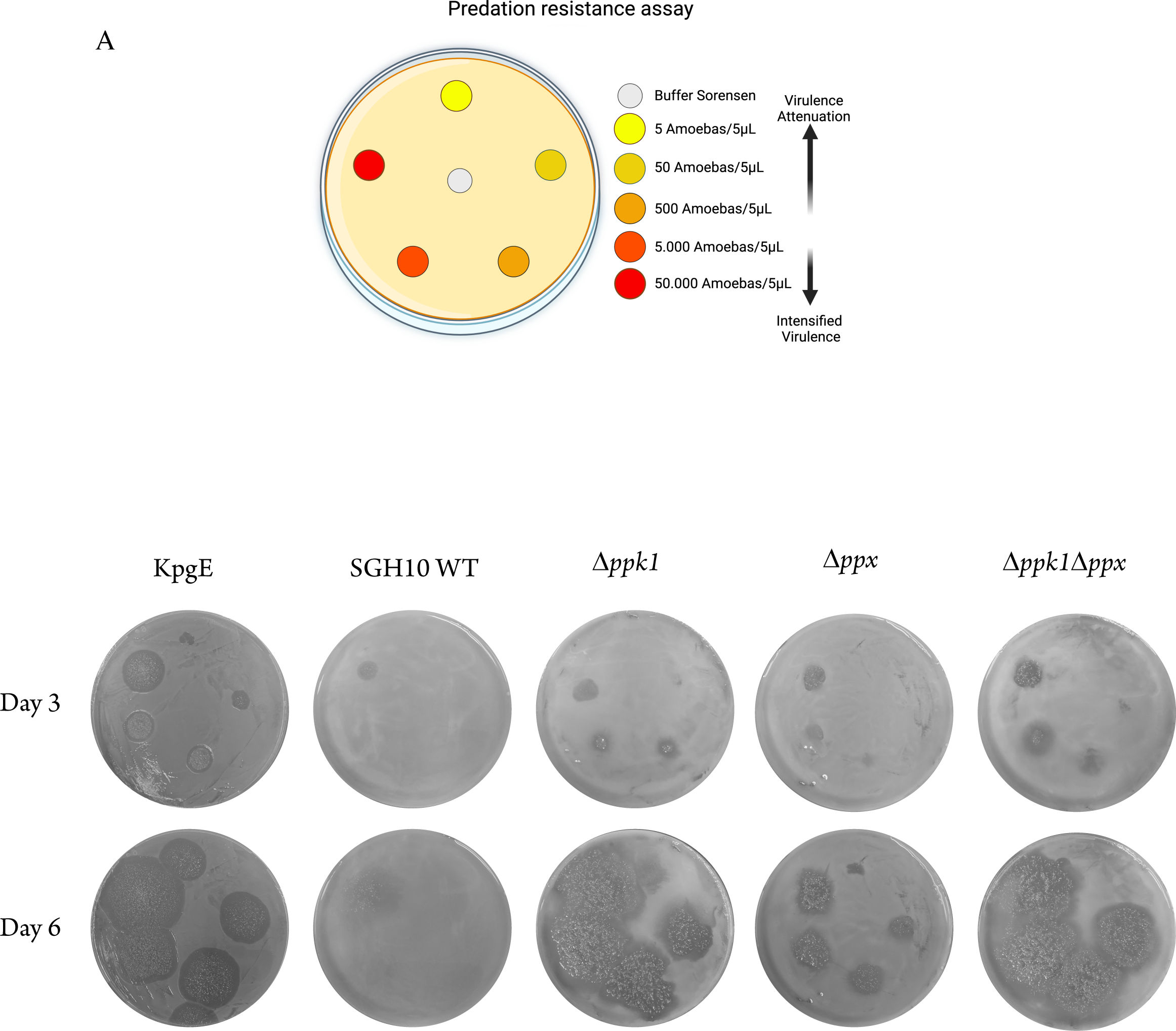
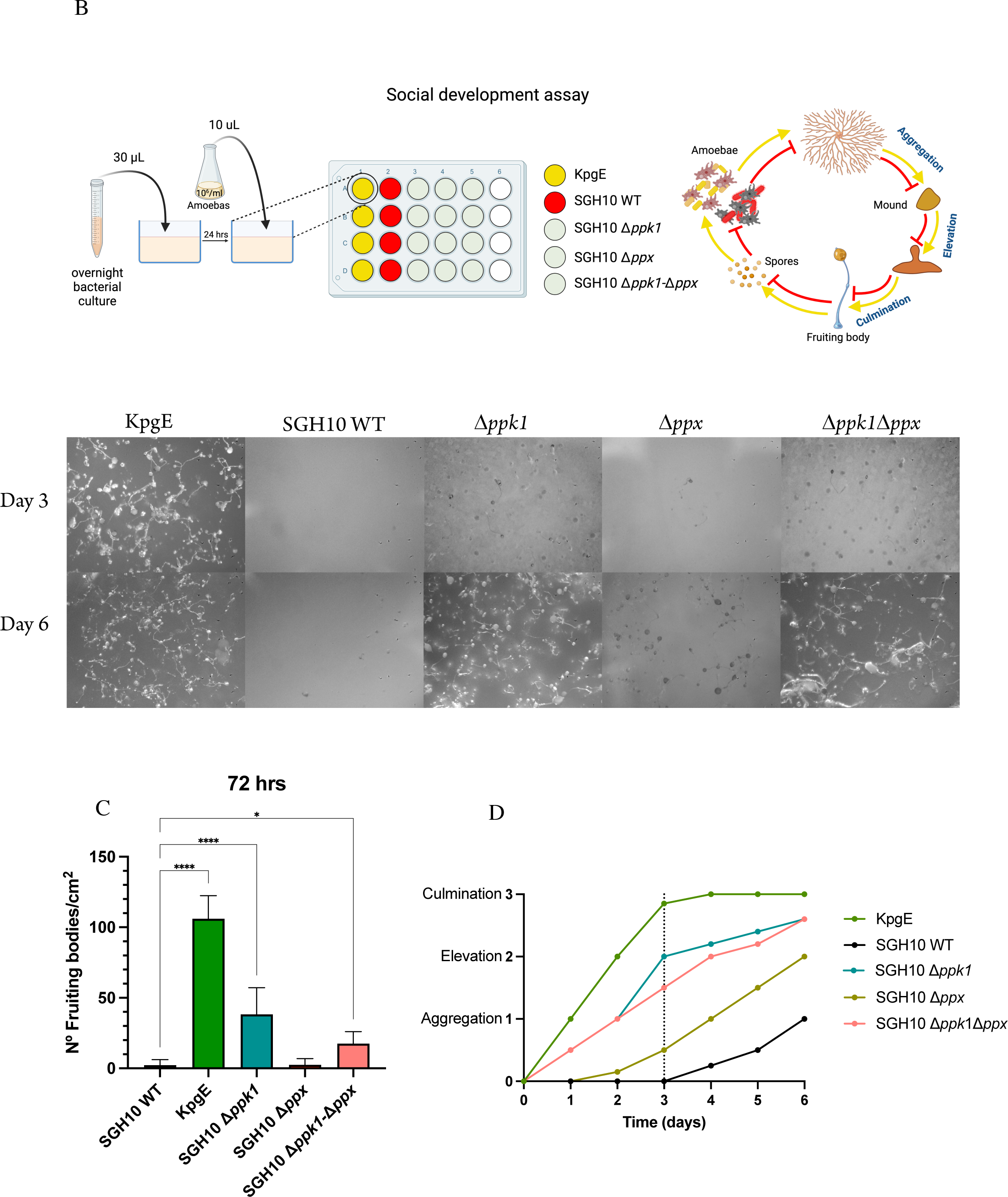
Biofilm formation is altered in polyP-deficient mutants. (A) Measurement/Quantification of relative biofilm formation in polyP metabolism mutants by 0.1% crystal violet staining. Overnight cultures were diluted to an OD600= 0.1 and plated in 96-well plates. The measurement was at 24 h at 595 nm. (B) EbbaBiolight 680 is an optotracer for facilitating biofilm visualization, specifically targeting cellulose and amyloid constituents within the biofilm matrix. Colonies were grown on agar plates under biofilm-forming conditions. EbbaBiolight 680™ was diluted in PBS 1:1000 and 100 µL were added into each well of a 96-well plate. Pick bacterial colonies from the agar plate and resuspend thoroughly into the pre-filled wells. Plates were placed in Plates in a spectrophotometer to quantify biofilm in the colony resuspensions. One-way ANOVA tests were performed for statistical analyses comparing the WT strain to the indicated mutants. *, P < 0.05 ****, P < 0.0001. (C) Confocal microscopy. SGH10 WT (1), SGH10 *Δppk1* (2), SGH10 *Δppx* (3), SGH10 *Δppk1-Δppx* (4) Overnight cultures were diluted 1:100 in LB and plated on a glass coverslip on a 6-well plate slanted at 80°. The plate was incubated for 24 hours. DAPI was used for biomass staining for 30 min, labeling DNA ex/em 358/462. Image obtained in LSM 710 Zeiss microscope, 20X. (D) Macrobiofilm. A macrobiofilm was formed by inoculating an overnight culture in LB broth onto LB agar plates supplemented with 40 μg/mL Thioflavin S. A 5 µL droplet of the culture was used for inoculation. The plates were subsequently incubated at 28 °C for 10 days.

### PPK1 is essential for HV *K. pneumoniae* virulence in *D. discoideum*

The findings from our study strongly indicate that PPK1 serves as the primary enzyme responsible for polyP synthesis in the *Klebsiella pneumoniae* SGH10 strain and that it is involved in the regulation of capsule and biofilm formation. The K1 capsular serotype, characterized by the trisaccharide composition of UDP-glucose, UDP-glucuronic acid and UGD-fucose ^40^, is recognized as a critical virulence factor that allows *K. pneumoniae* to evade immune system detection ^1^. The polysaccharide length has also been proposed to provide steric protection for important molecular patterns, such as lipid A from LPS ^2^. To investigate the impact of polyP and PPK1 deficiency on virulence, we used the social amoeba *D. discoideum* as a model host. When grown on lawns of avirulent bacterial strains, *D. discoideum* forms visible phagocytosis plaques. However, the amoeba fails to develop in the presence of virulent strains, and no phagocytosis plaques are observed ^41^. The versatility of this assay allows the determination of the minimum number of amoebas required to generate a phagocytosis plaque. Previous studies have shown that if phagocytosis plaques are formed with 500 or fewer *D. discoideum* cells, the bacteria exhibit susceptibility to predation, indicating attenuation of their virulence ^42^. We employed this assay to assess the impact of polyP metabolism on the virulence of the SGH10 strain. Results were recorded on the 3rd and 6th day of the assay (Fig. 4A). The KpgE strain was used as a control, and phagocytosis plaques were observed starting from day 3 at a concentration of 5 amoebas/5μL. For the SGH10 WT strain on the 3rd and 6th day, only one phagocytosis plaque was observed at a concentration of 50,000 amoebas/5μL, indicating the strain’s virulence towards amoebas. In the case of mutants in polyP metabolism (Δ*ppk1*; Δ*ppx*; Δ*ppk1*-Δ*ppx*), we observed an attenuation in virulence when compared to the SGH10 WT strain, as evidenced by the presence of phagocytosis plaques at a concentration of 500 amoebas/5μL. Furthermore, a differential impact on virulence attenuation was observed when comparing Δ*ppk1*, Δ*ppk1*-Δ*ppx* with Δ*ppx*. On the 6th day, large phagocytosis plaques were observed in the absence of the *ppk1* gene in both mutants, compared to the Δ*ppx* strain. This suggests that the lack of PPK1 has a more pronounced effect on attenuating virulence than the absence of PPX, despite both mutations leading to a decrease in virulence.

**Figure 4.**
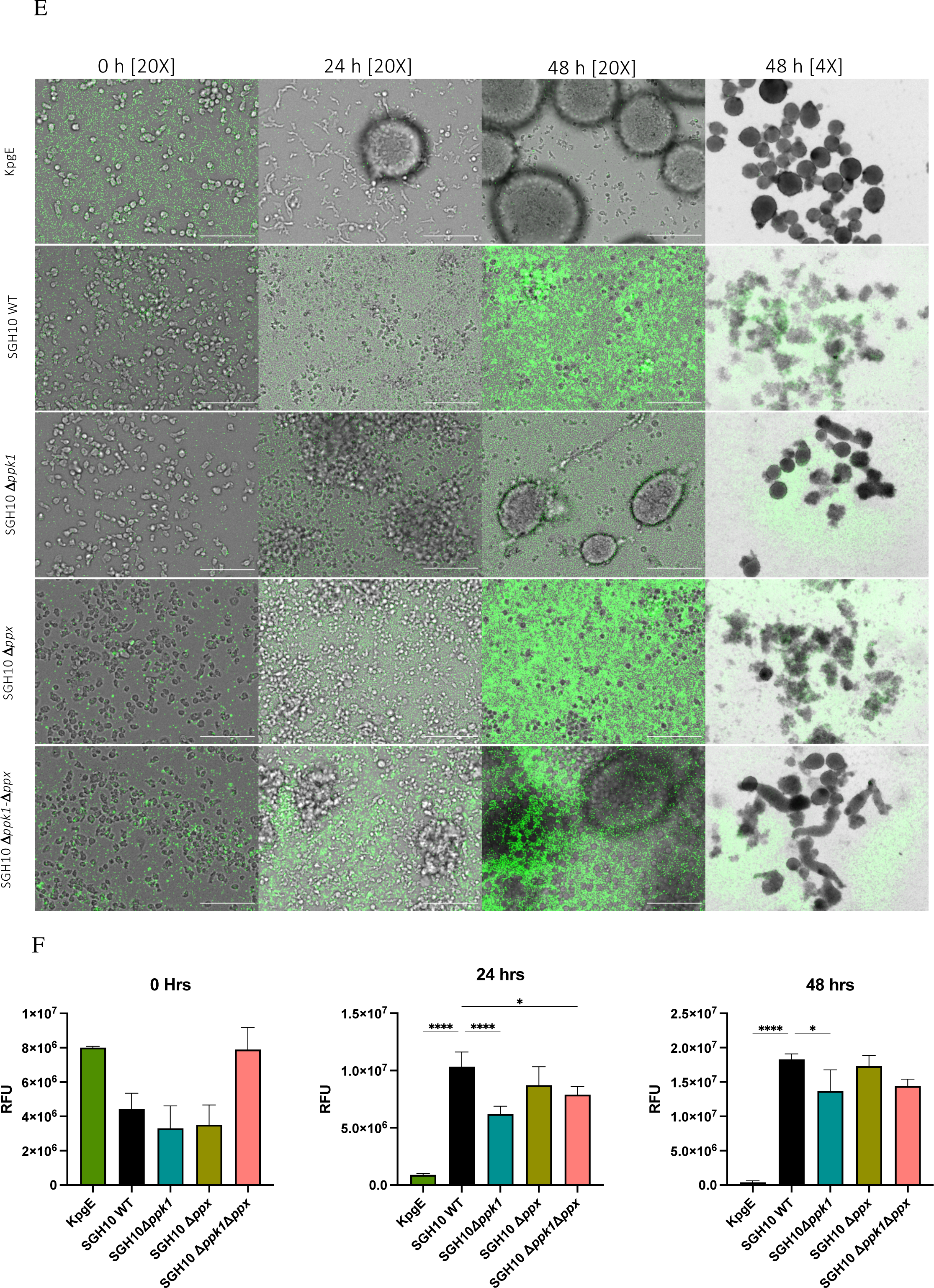
(A) Predation resistance assay. 300 µL of overnight culture were seeded onto a plate using a Digralsky loop to generate a bacterial lawn. The plate was left to dry and incubated for 24 hours at 23°C. *D. discoideum* was cultured in HL5 medium, and serial dilutions were prepared to obtain the following cell concentrations: 5 – 50 – 500 – 50.000 cells per 5 µL in Buffer Sorensen. Subsequently, 5 µL of *D. discoideum* serial dilutions were applied onto the bacterial agar plates, and the plates were allowed to dry and were incubated at 21°C for 6 days. Plaque formation was visually examined on days 3 and 6. Strains that did not enable amoebae growth were considered virulent for the amoeba. Bacterial strains that exhibited social development with 500 *D. discoideum* were considered sensitive to predation. (B) 30 mL of overnight cultures of the strains under study were inoculated in a 24-well plate on N agar to generate a lawn. They were incubated for 24 h at 23 °C. *D. discoideum* was adjusted to 10^6^/ml. A 10 mL drop of the dilution was inoculated on each law. Representative images of days 3 and 6 days are presented. The number of fruiting bodies was counted on the 3rd day and compared with the WT strain (C) and the social development of the amoeba was followed for six days (D). The KpgE commensal strain was used as a control. (E) Timelapse of a 48-hour phagocytosis assay in GFP channel (bacteria). KpgE was used as a control. Scale 100 μm. Images were acquired using an automated Lionheart FX microscope with a 20X objective. For multicellular development observation, a 4X objective was used. (F) Fluorescence quantification was performed in ImageJ with three biological replicates. One-way ANOVA tests were performed for statistical analyses comparing the WT strain to the indicated mutants. **** p < .00001, * p< 0.05.

Previous studies have demonstrated that pathogenic bacteria with high virulence capabilities can impede the social development of *D. discoideum* ^17,43,44^. Conversely, attenuated or avirulent bacteria have a minimal impact on the progression of social development in *D. discoideum*, which involves at least three stages: aggregation, including the formation of a phagocytosis plaque and subsequent cluster formation; elevation, involving the formation of slugs and fingers; and culmination, comprising the dispersal of fruiting bodies throughout the surface. The avirulent strain KpgE exhibited normal social development, culminating between days 2 and 3 with an approximate count of 110 fruiting bodies/cm2 on the third day (Fig. 4C-D). In contrast, the SGH10 WT strain displayed a significant delay in social development, failing to complete it within the 6-day study period, with only the aggregation phase observed on the sixth day (Fig. 4C). An average of 2.3 fruiting bodies/cm2 was observed on the third day (Fig. 4B-D). Mutants in polyP metabolism exhibited varying levels of virulence attenuation. Δ*ppk1* displayed the highest attenuation level, reaching a stage before the culmination of social development on the 6th day (Fig. 4D), with an average of 38 fruiting bodies/cm2 on the third day (Fig. 4C). This was followed by Δ*ppk1*-Δ*ppx*, which exhibited 17.5 fruiting bodies/cm2 on the third day, also reaching a stage before culmination (Fig. 4C). Δ*ppx* displayed lower levels of virulence attenuation, with a count of 2.5 fruiting bodies/cm2 on the third day (Fig. 4B-D).

To further substantiate our findings, we conducted epifluorescence microscopy to observe the phagocytosis *of D. discoideum* alongside bacteria constitutively expressing the green fluorescent protein GFP from the pBBR1-GFP-kan plasmid. Amoebas were infected with an MOI 10, and phagocytosis was monitored at 0-, 24-, and 48 hours post-infection (Fig. 4E). We noted a significant reduction in fluorescence intensity when the amoeba was infected with the avirulent strain KpgE from 0 to 24 hours (Fig. 4F). At 4X magnification, evident fruiting bodies were observed. Regarding the SGH10 WT strain, we observed a progressive increase in green fluorescence intensity from 0 to 48 hours (Fig. 4E). However, no fruiting bodies were observed across the well surface at 4X magnification; instead, we observed an aggregative behavior of amoebas. In the case of mutants in polyP metabolism, Δ*ppk1* exhibited significant levels of virulence attenuation. At 24 hours, clusters of amoebas were observed and by 48 hours, they had progressed in social development, with the presence of fingers, slugs, and fruiting bodies at 4X magnification (Fig. 4E). For the Δ*ppx* mutant, at 48 hours, no significant differences were observed compared to the WT strain regarding the increase in fluorescence intensity and the presence of amoeba aggregates (Fig. 4E-F). The Δ*ppk1*-Δ*ppx* mutant exhibited a similar pattern to the Δ*ppk* mutant, with amoeba aggregates at 24 hours and a lower increase in fluorescence intensity compared to the WT strain, along with the presence of fingers, slugs, and fruiting bodies at 48 hours. In conclusion, our study strongly supports the involvement of PPK1 in the virulence of hypervirulent *Klebsiella pneumoniae*.

### Global Proteomic Profiling of polyP metabolism mutants in hvKP

To investigate the underlying mechanisms responsible for the significant reduction in capsule production, biofilm formation, and virulence attenuation in the host model *D. discoideum,* we performed a shotgun proteomic analysis of mutants affecting polyP metabolism. Samples were obtained from colonies grown on LB agar at 37°C and subjected to LC-MS/MS analysis using a Data Dependent Acquisition method and label-free quantification (LFQ). The WT strain was compared to each mutant strain. Principal component analysis (PCA) on the identified protein dataset, as shown in Figure S4C, revealed the separation of the samples into their corresponding clusters like SGH10 WT, Δ*ppk1*, Δ*ppx*, and Δ*ppk1*-Δ*ppx* groups. Later, among the 2,287 quantifiable proteins identified, 302, 728, and 673 were determined to be Differential Expression Proteins (DEPs) in the comparisons of WT vs. Δ*ppk1*, WT vs. Δ*ppx*, and WT vs. Δ*ppk1*-Δ*ppx*, respectively (Table S2). The quantifiable proteins resulting from these comparisons were visualized as volcano plots (Fig. 5A), where significantly upregulated proteins were represented in blue and downregulated proteins in red. To further investigate potential virulence factors associated with the observed results, the differentially expressed proteins were compared against the VFDB database and those with a ≥50% identity were selected for further analysis. The virulence factors were categorized into anti-phagocytosis, adherence, iron uptake, and toxins and visualized in a heatmap (Fig. 5B), a network of curated interactions (Fig. S5) and a chordplot (Fig. S6). Regarding adherence, we observed a significant reduction in essential proteins associated with capsule synthesis, transport, and regulation in *K. pneumoniae*, particularly in the double mutant Δ*ppk1*-Δ*ppx.* This mutant exhibited decreased expression of GalF, Wzi, Wza, WcsT, Wca, Ugd, and Gmd compared to the wild-type strain. Furthermore, in Δ*ppk1*, Wzi and Ugd proteins were downregulated, while Δ*ppx* showed an overexpression of Wzi compared to the SGH10 WT strain. Concerning iron uptake, proteins related to the synthesis of siderophores such as aerobactin, enterobactin, salmochelin, and yersiniabactin were found to be overexpressed in all polyP mutants, except for yersiniabactin in Δ*ppk1*, where no significant differences were observed compared to the wild-type strain. Both Δ*ppx* and Δ*ppk*1-Δ*ppx* exhibited upregulation of operons involved in the synthesis of aerobactin and enterobactin, as well as the protein YbtS, which is implicated in the final step of siderophore union and modification, playing a crucial role in the overall siderophore biosynthesis process. In Δ*ppk1*, several proteins involved in siderophore synthesis, including IucA (aerobactin), EntB, EntC, FepC (enterobactin), and IroD (salmochelin), were found to be overexpressed. Regarding the toxin category, in all polyP metabolism mutants, a significant downregulation was observed in most of the proteins from the colibactin genotoxin biosynthetic cluster. This downregulation was particularly evident in Δ*ppx* and Δ*ppk1*-Δ*ppx*, affecting proteins such as ClbB, ClbD, ClbF, ClbG, ClbH, ClbI, ClbK, ClbL, ClbN, and ClbQ. In the case of Δ*ppk1*, the main protein involved in colibactin biosynthesis, ClbB, was downregulated along with ClbD. In the Lipid A category, we observed a significant increase in proteins involved in the biosynthesis and modification of Lipid A, such as ArnA, arnB, ArnC, PagP, and LpxO in Δ*ppx* and Δ*ppk1*-Δ*ppx*. Δ*ppk1* displayed similar differential expression levels in these proteins; however, no significant differences were observed for ArnA. Additionally, proteins PagP, LpxC, and FabR, related to Lipopolysaccharide (LPS) biosynthesis and modification, exhibited decreased expression levels compared to the WT strain. This prompted us to evaluate whether the differential expression of proteins involved in Lipid A modification would confer colistin resistance (Fig. S7). The results did not show increased colistin resistance, with a 4 µg/ml MIC. Finally, in Figure 5C, we present a z-score plot that considers the abundances of 38 proteins related to each virulence category (adherence, anti-phagocytosis, toxin, siderophores, lipid A). The analysis reveals that the Δ*ppk1* mutant exhibited a negative z-score compared to the WT for all virulence factors reported in the VFDB database. In conclusion, our global proteomic profiling of polyP metabolism mutants revealed significant alterations in protein expression related to biofilm formation, capsule regulation, iron uptake, and toxin production, corresponding to key virulence-associated traits in hvKP.

**Figure 5.**
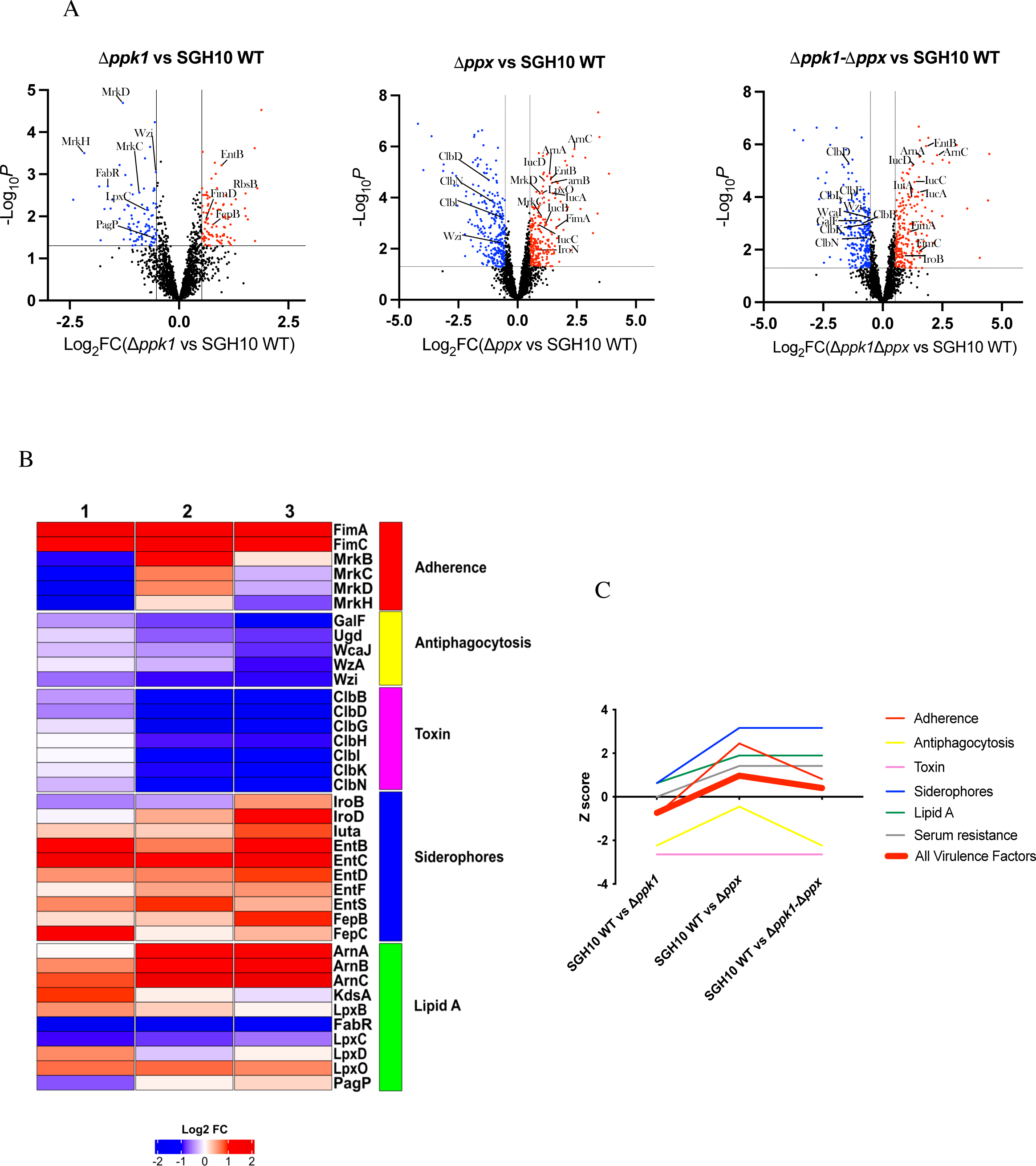
Global proteomic profiling of polyP metabolism mutants in hvKP. (A) Volcano plot of differentially expressed proteins from each polyP metabolism mutant strain. These colored points indicate different proteins that display large magnitude fold-changes (x-axis) and high statistical significance (-log10 of p values, y-axis). The dashed horizontal line shows the cut-off p-values, and the two vertical dashed lines indicate down (blue) /up (red) differentially expressed proteins. Black points in the significant region mean these proteins do not satisfy these conditions. (B) Heatmap shows differentially expressed virulence factors (VF) of polyP metabolism mutants in hvKP. The proteome was compared with the VDFB database, and the VFs were classified according to the categories: Adherence, anti-phagocytosis, siderophores, toxin, lipid A, serum resistance, and secretion system. (C) Z-score plot illustrating the global relative abundance of each of the proteins associated with virulence factors. Positive z-scores indicate proteins with higher abundance, while negative z-scores indicate proteins with lower abundance compared to the overall mean. The red line represents the average of all z-scores for the virulence factors.

**Figure 6.**
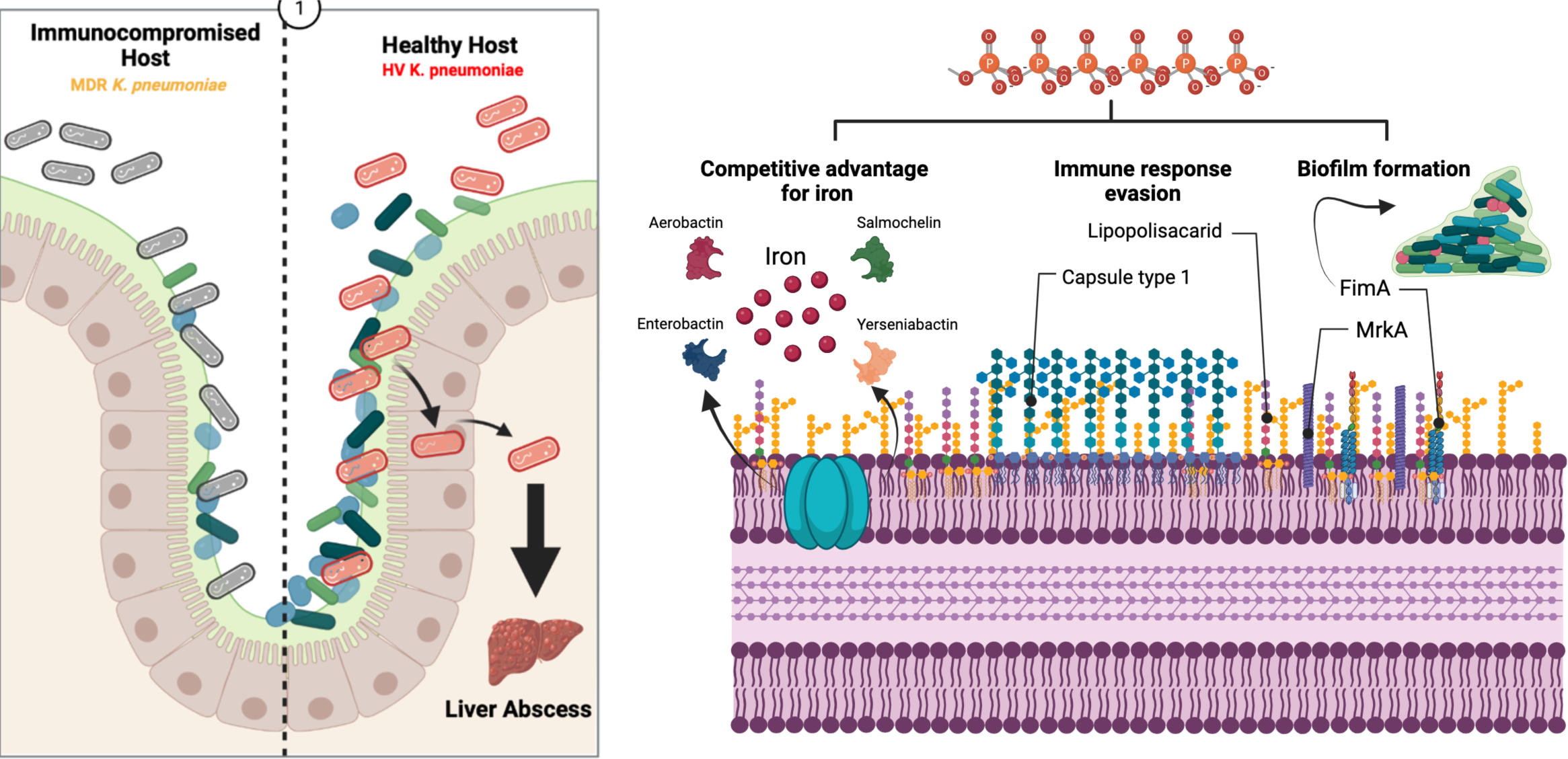
Model. In classic *Klebsiella pneumoniae* strains causing nosocomial infections, colonization of the intestines typically requires prior dysbiosis induced by factors like antibiotics or immunocompromised patients. In contrast, hypervirulent *Klebsiella pneumoniae* strains could colonize the intestines and invade organs such as the liver, leading to abscesses throughout the body, regardless of the patient’s prior condition. To achieve this, they possess various virulence factors that facilitate colonization, invasion, and dissemination from the site of infection. In our study, we established a connection between polyP metabolism and critical aspects of the expression and regulation of key virulence factors. This included alterations in capsule formation, the hypermucoviscous phenotype, decreased biofilm formation, and changes in virulence.

## DISCUSSION

1. *K. pneumoniae* is a growing health threat due to its antibiotic resistance and the emergence of highly virulent strains. These strains can infect healthy individuals and are adept at causing severe infections, posing risks to hospitalized patients and the general population^45^. Traditional approaches targeting bacterial viability through antibiotic use have led to the selection of multidrug-resistant strains, necessitating a shift in strategy. In line with the principle of “Disarm and not kill,” there is an increasing recognition that targeting virulence mechanisms rather than bacterial viability could provide a promising alternative to mitigate the risk of resistance development. In this study, we aimed to investigate the involvement of polyP metabolism in the hypervirulent ST23 *Klebsiella pneumoniae* SGH10 strain, specifically focusing on the enzyme PPK1 and its implications on capsule formation, biofilm formation, and virulence ^46–48^.

In the hypervirulent strain SGH10, we observed that PPK1 is the primary enzyme responsible for synthesizing inorganic polyphosphates. As anticipated, the absence of PPK1 and a marked reduction in polyP did not substantially impact viability under favorable growth conditions ^28^. However, when exposed to minimal conditions (stress response), a 2-h delay in reaching exponential growth was observed. This delay is expected, as nutrient scarcity is known to be one of the main triggers for polyphosphate synthesis, and its absence leads to a noticeable growth delay that eventually recovers when increasing concentrations of polyP were added.^16^. In the PPX mutant, a drastic increase in polyP formation or accumulation in response to nutritional downshift in the MOPS medium (0.4% glucose, 0.1 mM K2HPO4) was not observed. The classical model proposed by A. Kornberg’s group for *E. coli* suggests that a stress response such as amino acid starvation leads to the accumulation of (p)ppGpp, which interacts with PPX and inhibits it, promoting polyP accumulation. However, based on recent evidence, other factors such as DksA and GreA proteins significantly regulate polyP accumulation in response to amino acid starvation ^49,50^. This could explain our observations in the Δ*ppx* mutant. However, there is no pronounced effect on viability, suggesting that PPK1 could be an attractive target for antivirulence strategies in this hypervirulent strain. Similar results were reported for *P. aeruginosa* PAO1, where some PPK1 inhibitors affected its virulence, resembling Δ*ppk1* knockout mutant phenotype^51^.

Several previous studies have demonstrated that the K1 capsule present in the SGH10 strain is a crucial virulence factor that enables evasion of immune recognition and resistance to phagocytosis ^34^. Recently, it has been shown that the length of the capsular polysaccharide is relevant to the characteristic HMV phenotype of hypervirulent strains, although not exclusive to them ^5,6^. This length regulation is mediated by the RmpD protein, which interacts with Wzc to regulate polysaccharide length. Our results indicate that PPK1 promotes the HMV phenotype and capsule expression or synthesis. The Δ*ppk1* mutant displayed longer sedimentation times during low-speed centrifugation, a standard method to assess mucoviscosity ^52^. Additionally, both the Δ*ppk1* and double mutant cells exhibited lower capsule levels, indicated by reduced uronic acid content and an altered distribution of capsule polysaccharides, which is distinct from the SGH10 strain.^34^. In addition to the findings in *Klebsiella pneumoniae*, a Δ*ppk1* mutant of *P. aeruginosa* has also shown an altered surface structures ^20^. In *Neisseria meningitidis*, it has been demonstrated that polyP can interact with the membrane, forming a capsule-like structure ^53^. This indicates that the role of PPK1 and polyP in capsule synthesis and structure may be conserved across different bacterial species.

The presence of polyP appears crucial for the proper assembly and organization of the capsule, and its absence or disruption can lead to abnormalities in the superficial structures. The defective membrane pattern observed in the Δ*ppk1* mutant of *P. aeruginosa* suggests that polyP regulates external biomolecule biosynthesis or assembly. Proteomic analysis of the Δ*ppk1* mutant and Δ*ppk1*-Δ*ppx* revealed decreased expression of proteins involved in capsular polysaccharide synthesis, such as Wzi-Ugd (Δ*ppk1*), as well as GalF, GalU, Wza, WcaJ, Ugd (Δ*ppk1*-Δ*ppx*). This downregulation could explain the decrease in mucoviscosity, as reported in the HV *K. pneumoniae* strain NTUH-K2044 ^10^. These findings suggest that polyP metabolism may be relevant in regulating capsule expression and the HMV phenotype.

Inorganic polyP may act as a signal molecule, triggering specific regulatory pathways controlling capsule biosynthesis. Similarly to *E. coli* and *P. aeruginosa*, where PPK1 is predominantly associated with the membrane ^37^, it is possible that in *K. pneumoniae*, this preferential localization has regulatory roles in biofilm formation and capsule and membrane processes. Furthermore, polyP may act as a phosphate donor to form sugar linkages in the capsule synthesis pathway ^54^. Phosphate is crucial in numerous biological molecules and processes within bacteria, encompassing the synthesis of polysaccharides integral to the bacterial capsule. Mutants deficient in PPK1 and PPK1-PPX have previously exhibited a phenotype associated with phosphate-related effects in *E. coli* ^55^. Phosphates are commonly present as phosphate groups in the structure of carbohydrates and other biomolecules. Polysaccharides, which constitute the capsule, can contain phosphate groups as part of their chemical structure or as a modification of their structure ^56^. These phosphate groups may correspond to side chains or substitutions on the sugar molecules forming the polysaccharide backbone ^57^. The presence of phosphate groups in the capsule can have functional implications, contributing to the overall charge and hydrophilicity of the capsule and then influencing its interactions with the environment, such as host tissues and immune cells. Phosphate groups can also be involved in the binding of cations and other molecules, further modulating the properties and functions of the capsule.

Biofilm formation is the primary lifestyle of bacteria in their natural state and serves as a successful strategy to thrive in niches subjected to various stressors and the most efficient way for the trafficking of mobile genetic elements ^58^. There is a broad consensus in the literature regarding the ability of both classical and multidrug-resistant Klebsiella pneumoniae strains to form biofilms in various niches, particularly in immunocompromised individuals ^59–62^. The *fim* and *mrk* operons, responsible for encoding type I and type III fimbriaes, respectively, play a crucial role in the initial adhesion to both biotic and abiotic surfaces ^63^. Regarding HV strains, it has been observed that their elongated capsular polysaccharides can cover the fimbriae, hindering their contact with surfaces and resulting in weaker biofilm formation ^9,64^. However, conflicting reports suggest that the capsule may promote the formation of biofilm structures, particularly within infectious contexts ^10,11,65^. Our findings demonstrate the capability of the SGH10 strain to form biofilms on polystyrene plates, glass, and at the air-liquid interface. The Δ*ppk1* and Δ*ppk1*-Δ*ppx* mutants exhibited reduced biofilm formation compared to the WT strain. These results are consistent with previous reports in various models ^28,66–69^. Interestingly, the use of mesalamine, a drug used in the treatment of ulcerative colitis that also inhibits PPK1 ^30^, showed similar levels of biomass reduction in the biofilm as observed in Δ*ppk1* and Δ*ppk1*-Δ*ppx* mutant cells, confirming the close association between biofilm formation and polyphosphates.

The proteomic analysis further confirms the relevance of polyP metabolism in biofilm formation. In the case of the Δ*ppk1* mutant, we observed a decrease in the transcription levels of *mrkC*, *mrkD*, and *mrkH* genes, accompanied by an increase in *fimA* and *fimD*. Conversely, the Δ*ppx* mutant significantly increased Fim (FimA, FimC, FimD) and Mrk (MrkB, MrkC, MrkD) proteins. Similarly, the Δ*ppk1*-Δ*ppx* mutant exhibited elevated levels of FimA and FimC proteins. This dysregulation can be partially explained by a previous report pointing out that polyP degradation during the stationary phase promotes biofilm formation ^70^. Consequently, the insufficient levels of polyphosphates in the Δ*ppk1* and Δ*ppk1*- Δ*ppx* mutants lead to the downregulation of proteins from the *mrk* operon. However, this explanation does not fully account for the increased proteins from the *fim* operon, indicating the involvement of broader regulatory mechanisms. A recent study has reported a direct correlation between biofilm formation and the colibactin operon in isolates of *E. coli* and *K. pneumoniae* ^71,72^. Furthermore, since it has been demonstrated that the inactivation of PPK by mutagenesis reduces the promoter activity of *clbB* (the gene that encodes for an essential enzyme for colistin biosynthesis) and decreases the production level of colibactin ^73^, it is possible that the absence of this enzyme also leads to a decrease in biofilm formation through this pathway. Recently, a phenotypic switch between type III fimbriae and the hypermucoviscous capsule in response to iron levels has been reported in the SGH10 strain.^70,27^ The iron-regulated outer membrane protein IroP, encoded in the virulence plasmid, inhibits the expression of the *mrk* operon under low iron conditions, promoting biofilm formation. Since polyphosphates (polyP) are directly involved in intracellular iron homeostasis ^74^, the observed alterations in the capsule and biofilm might be related to this recently described phenomenon.

In addition, the role of polyP in forming functional amyloids in prokaryotes and its potential relationship with the appearance of these fibers in the context of biofilms should be considered ^75^. Several amyloid-like proteins, such as curli in *E. coli*, have been described, highlighting the potential significance of amyloid formation in bacterial biofilms. Although *K. pneumoniae* lacks the curli operon, the GIE492 genomic island enabling the production of the antimicrobial peptide microcin E492 is highly prevalent among hvKP ^76^. This microcin forms amyloid fibers in the culture supernatant of producing cells^77^, although the meaning of this phenomenon is still unclear. Possibly, it could be a structural part of biofilms. Furthermore, it is worth considering the potential role of interbacterial communication and quorum sensing in biofilm formation. The coordinated behavior of bacteria within the biofilm community relies on intricate signaling networks that enable collective responses and establish structured microbial communities ^78^. Further investigations into the interplay between polyphosphates, quorum sensing systems, and other signaling pathways will provide valuable insights into the complex dynamics of biofilm formation and the development of targeted interventions. Based on our findings, we propose that polyphosphates could serve as potential extracellular signaling molecules within the biofilm matrix, acting as an energy source for dynamic biofilm processes and as a structural component due to their polyanionic nature, like eDNA ^54^. Further studies are warranted to ascertain the essential role of polyphosphates as constituents of the biofilm rather than solely intracellular signaling molecules. Understanding the intricate regulatory pathways involved, including interbacterial communication and quorum sensing, will provide valuable insights for developing strategies to control biofilm-associated infections and enhance the efficacy of antimicrobial therapies.

Unlike classical cKP strains, which the host defense system can rapidly clear during infection, hvKP strains can evade and resist macrophage- and neutrophil-mediated killing, generally considered an extracellular pathogen ^2^. However, reports show that hvKP strains can persist intracellularly for more than 24 hours by diverting the canonical endocytic pathway and surviving in a phagosome not associated with lysosomes ^79^. *D. discoideum* has been successfully used as a host model to study virulence and phagocytosis resistance ^43,80^. Here, the predation resistance assay demonstrated the contribution of polyP metabolism to the virulence of the hypervirulent SGH10 strain (Fig. 5A). This reflects the involvement of PPK1 in mediating key aspects of virulence in the SGH10 strain. On the other hand, the amoeba model allows us to evaluate alterations in social development, which, under standard laboratory conditions, takes between two and three days to go through the stages of aggregation, elevation, and culmination ^17^. In the presence of a virulent strain, this process can be delayed or may not occur at all. Δ*ppk1* and Δ*ppk1*-Δ*ppx* mutants showed reduced virulence with more fruiting bodies by the 3rd day. These results provide insight into the influence of polyP metabolism in this hypervirulent strain, as documented in other pathogens. The significant decrease in capsule formation, the HMV phenotype, and a reduction in colibactin expression could be responsible for the attenuation of virulence in the studied model. Furthermore, long-chain bacterial polyphosphates (> 300 mer) have been shown to interfere with immune cell signaling pathways, preventing the arrival of macrophages and neutrophils at the site of infection ^25^ and inhibiting phagolysosome formation in *D. discoideum* ^24^. The lack of polyP could make the bacteria more vulnerable to the immune response, as it cannot manipulate or prevent phagolysosome fusion.

We were particularly intrigued by the significant increase in the pool of siderophores in all mutants, especially in Δ*ppx* and Δ*ppk1*-Δ*ppx* mutants. It has been recently reported that polyphosphates (polyP) can sequester free iron, acting as a reservoir under non-stress conditions and blocking the formation of reactive oxygen species (ROS) during stress by inhibiting the Fenton reaction ^74^. We believe that alterations in polyP metabolism might dysregulate the expression of siderophores under non-stress conditions, particularly in the Δ*ppx* mutant that accumulates slightly higher levels of polyP. The absence of PPX implies a reduced capacity for polyP degradation and consequently, a diminished release of intracellular iron under these conditions. This could serve as a signal to induce the expression of siderophores to compensate for the lack of free iron necessary for metabolic activities. Although it has been documented that aerobactin is essential for the pathogenicity of the strain under study ^7^, it has also been reported that the K1 and K2 capsule types have greater relevance in virulence ^81,82^.

Despite the extensive body of literature focusing on PPK1, the role of PPX in the physiology of either non-pathogenic or pathogenic bacteria remains largely unexplored. Particularly regarding the virulence traits manifested in Δ*ppx* mutants in bacteria, it has been reported that PPX is essential for the pathogenesis of *Mycobacterium tuberculosis* ^83^, Bacillus cereus ^84^ and Neisseria meningitidis ^85^. Our findings in Δ*ppx* mutant demonstrated no significant alteration in its ability to evade phagocytosis and impact the amoeba’s social development.

In summary, our research highlights the role of polyP synthesis in modulating extracellular biomolecules and virulence factors in hypervirulent ST23 *K. pneumoniae*. The study offers valuable insights into possible therapeutic targets for its treatment. However, more research is needed to fully understand the intricate relationship between polyP metabolism, siderophores, and the virulence of the ST23 *Klebsiella* hypervirulent strains.

## CONCLUSION

This study investigated the impact of polyphosphate metabolism in the hypervirulent strain of *Klebsiella pneumoniae* SGH10. It was found that the enzyme PPK1 is essential for polyphosphate synthesis and plays a role in capsule formation, hypermucoviscosity, and biofilm formation. The absence of PPK1 affected growth under stress and the expression of genes related to capsules and biofilms. Furthermore, the mutants showed changes in siderophore production, suggesting an interaction with iron homeostasis. These findings highlight PPK1 as a potential target to mitigate virulence and enhance antimicrobial strategies in hypervirulent strains of *K. pneumoniae*.

## METHODS

### 1. Strains, plasmids, and growth conditions

The details of the used strains and plasmids can be found in Table S2. *E. coli* and *K. pneumoniae* strains were grown routinely at 37°C in LB broth (10 g/L Tryptone, 5 g/L Yeast Extract, 5 g/L NaCl). For solid medium, 1.5% w/v agar was added. MOPS medium supplemented with 0.4% glucose and 0.1% K2HPO4 was used for the nutrient deprivation assay. For the macrobiofilm assays, LB agar without NaCl supplemented with Thioflavin S (40 μg/mL) or Congo Red (40 μg/mL) was used, and 100 μg/mL Ampicillin or Carbenicillin, 50 μg/mL Kanamycin, or 10 μg/mL Tetracycline was used for antibiotic selection.

### 2. Growth conditions of *D. discoideum*

We obtained the *D. discoideum* strain AX4 (DBS0302402) from the Dicty Stock Center ^86–88^ and cultured it using standard protocols ^89^. Briefly, the *D. discoideum* vegetative cells were maintained at 23°C in SM medium containing 10 g/L glucose, 10 g/L peptone, 1 g/L yeast extract, 1 g/L MgSO4 × 7H2 O, 1.9 g/L KH2PO4, 0.6 g/L K2HPO4, and 20 g/L agar. They were grown on a confluent lawn of *Klebsiella aerogenes* DBS0305928 (KpgE, also from the Dicty Stock Center). Before the assays, amoebae were grown at 23°C with agitation (180 rpm) in liquid HL5 medium, which contained 14 g/L tryptone, 7 g/L yeast extract, 0.35 g/L Na2HPO4, 1.2 g/L KH2 PO4, and 14 g/L glucose at pH 6.3. These cultures were axenic, meaning they were free of bacteria. We harvested amoebae in the early exponential phase (1-2 × 10^6^ cells/mL) and centrifuged them at 2000 rpm for 5 min. Before the infection assays, we discarded the supernatant. We washed the pellet three times using Soerensen buffer (2 g/L KH2 PO4, 0.36 g/L Na2HPO^4^ × 2 H2O, pH 6.0) or adjusted it to 10^6^ cells/mL in HL5 medium for social development assays. We determined the number of viable amoeba cells by performing Trypan blue exclusion and counting in a Neubauer chamber. For the social development assay involving *D. discoideum*, Agar N (1 g peptone, 1 g glucose, 20 g agar in 1 L of 17mM Soerensen phosphate buffer) was employed as the substrate. Meanwhile, Agar SM served as the substrate in the predation resistance assay.

### 3. Construction of *K. pneumoniae* SGH10 mutants

For *K. pneumoniae* scarless site-directed mutagenesis, we followed the protocol described previously^34^. Briefly, we first amplified ∼1000 bp fragments upstream and downstream of the target gene using *K. pneumoniae* SGH10 genomic DNAas template and the Q5 high-fidelity DNA polymerase (New England Biolabs). Then, these fragments were assembled along with the previously PCR-amplified vector pR6KTet-SacB using the NEBuilder® HiFi DNA Assembly Master Mix (New England Biolabs). The assembled constructs were then transformed by electroporation into *E. coli* S17-1λpir (donor strain), and then conjugated into *K. pneumoniae* SGH10 (recipient strain), selecting for transconjugants by plating in LB agar plates supplemented with tetracycline (50 µg/mL) and ampicillin (100 µg/mL). Afterward, several rounds of sucrose counterselection were done to select for the second recombination event, leading to the loss of the plasmid backbone comprising the *sacB* gene. For this, single tetracycline-resistant crosses were passaged in an LB medium without sodium chloride but supplemented with 20% sucrose. Successful deletion of the gene of interest was evaluated by PCR amplification of the respective region from tetracycline-sensitive double recombinant clones and confirmed by Sanger sequencing. Moreover, to rule out possible off-target mutations, we performed Illumina whole-genome sequencing of the mutant strains, and the reads were mapped to the *K. pneumoniae* SGH10 complete genome using BWA-MEM ^90^.

### 4. Quantitative proteomics profiling

The samples were obtained by selecting colonies grown under ideal conditions directly from an LB-Agar culture plate at 37°C. These were then frozen for processing.

#### a. Protein Extraction

Protease/phosphatase inhibitor (#1861284, Thermo Scientific) was added to each sample at a 1X final concentration. Then, the samples were lyophilized and resuspended in 8M urea with 25 mM ammonium bicarbonate pH 8 and later, they were homogenized using ultrasound for 1 min with 10 s pulses (on/off) at an amplitude of 50% using a cold bath. Then, they were incubated on ice for 5 min and later centrifuged to remove debris at 19,000 x g for 10 min at 4°C. Samples were immediately quantified using the Qubit Protein Assay reagent (#Q33212, Invitrogen).

#### b. MS Preparation

The resulting proteins were subjected to chloroform/methanol extraction. Then, they were equilibrated at room temperature for 10 min, centrifuged at 15,000 x g for 5 min at 4°C, and discarded into the supernatant. The resulting pellet was washed 3 times with cold 80% acetone. Subsequently, the protein pellet was dried in a rotary concentrator. Samples were resuspended in 30 μL 8M Urea and 25 mM ammonium bicarbonate. Then, they were reduced with DTT to a final concentration of 20 mM in 25 mM ammonium bicarbonate and incubated for 1 hour at room temperature. They were then alkylated by adding iodoacetamide to a final concentration of 20 mM in 25 mM ammonium bicarbonate and incubated for 1 hour in the dark at room temperature. Subsequently, the samples are diluted 8 times with 25 mM ammonium bicarbonate.

Digestion was performed with sequencing grade Trypsin (#V5071, Promega) in a 1:50 ratio of protease: protein (mass/mass) and incubated for 16 h at 37°C. Digestion was stopped by pH adding 10% formic acid. Then, the samples were subjected to Clean Up Sep-Pak C18 Spin Columns (Waters), according to the supplier’s instructions. Subsequently, the clean peptides were dried in a rotary concentrator at 2000 rpm overnight at 40°C.

#### c. LC-MS/MS (Liquid Chromatography – Tandem Mass Spectrometry)

200 ng of the tryptic peptides obtained in the previous step were injected into a nanoELUTE nanoUHPLC (Bruker Daltonics) coupled to a timsTOF Pro mass spectrometer (“Trapped Ion Mobility Spectrometry – Quadrupole Time Of Flight Mass Spectrometer”, Bruker Daltonics) using an Aurora Ultimate column. UHPLC (25 cm x 75 μm ID, 1.6 μm C18, CSI, IonOpticks, Australia). Liquid chromatography used a 90-min gradient of 2% to 35% buffer B (0.1% Formic Ac. - Acetonitrile). The collection of results was performed using the TimsControl 2.0 software (Bruker Daltonics) under 10 PASEF cycles, with a mass range of 100-1,700 m/z, capillary ionization of 1,500 V, and a temperature of 180°C, TOF frequency of 10 KHz at a resolution of 40,000 FWHM.

#### d. Protein Identification

The data obtained were analyzed with the MSFragger 3.5 software (Kong et al., 2017) through the Fragpipe v18.0 platform (https://fragpipe.nesvilab.org/) using the “default” workflow in a data analysis server consisting of 48 cores and 512 Gb of RAM. Mass tolerance parameters of 50 ppm were used, using monoisotopic masses and 0.05 Da fragment ions. Among the digestion options, trypsin was used as the enzyme-specific digestion mode and a maximum of 2 missed cleavages per peptide. The following were used as post-translational modifications (PTM): Cysteine carbamidomethylation, as fixed PTM: Methionine oxidation (M), N-terminal acetylation, Asparagine and Glutamine (NQ) deamination, as variable PTMs. The database used for identification was the proteome of *Klebsiella pneumoniae* strain SGH10 (5,405 total entries). The FDR estimate was included using a decoy database. As a filter, an FDR

≥ 1% and 1 minimum unique peptide per protein were used for identification.

#### g. Protein Quantification LFQ (Free Label Quantification)

The columns corresponding to the Uniprot access codes and intensity values were selected using the protein identification results. Data were concatenated, and missing values in the intensity columns were imputed. The resulting matrix was subjected to normalization by medians. Differential expression proteins were determined by applying a T-Test with a Benjamini-Horschberg multiple correction test (adjusted p-value < 0.05). DEPs (Differentially Expressed Proteins) were identified between the comparisons: *Δppk1*vs. WT, *Δppx* vs. WT and *Δppk1*-*Δppx* vs. WT.

### 5. Calculation of the z-score

The z-score was calculated based on the logFC of each protein belonging to the virulence categories derived from VFDB (Adherence, antiphagocytosis, toxin, siderophores, lipid A, serum resistance). It’s important to note that this z-score doesn’t refer to the standard score typically used in statistics but is a simple calculation aimed at indicating whether the biological process, molecular function, or cellular component is more likely to decrease (resulting in a negative value) or increase (resulting in a positive value). The calculation was performed using the following formula:

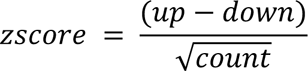

The z-score was determined based on the number of proteins assigned as upregulated (logFC>0) or downregulated (logFC<0), denoted as “up” and “down,” respectively, in the dataset. Subsequently, the average z-scores of each category were calculated for each mutant and plotted as “All virulence factors.”

### 6. Growth and polyP production under nutrient deprivation

Mutant strains with alterations in polyP metabolism and the SGH10 WT strain were cultured overnight. A 1:1000 dilution in LB or MOPS medium was inoculated into a 96-well plate (the MOPS medium was supplemented with 0,4% glucose and 0.1 mM K2HPO4). The plate was then incubated for 24 h in a TECAN infinite 200 pro plate reader at 37°C and 180 rpm. Readings were taken every 10 min (n = 6) to measure the optical density at 600 nm (OD600).

To investigate whether PPK1 deficiency affects bacterial polyP accumulation, a polyP quantification assay was performed ^91^. Briefly, cultures of *K. pneumoniae* SGH10 and polyP mutants grown overnight were diluted into fresh LB broth and cultured at 180 rpm and 37°C for 2 h. Then, 1 mL of bacterial cells was centrifuged twice in MOPS low-phosphate minimal medium (containing 0.4% glucose and 0.1 mM potassium phosphate) and incubated at 37°C for 2 h to stimulate polyP accumulation. Then, cells were harvested by centrifugation in a 1,5 mL. The supernatant was entirely removed from the cell pellets. Subsequently, the cell pellets were resuspended in 250 μL of GITC lysis buffer (consisting of 4 M guanidine isothiocyanate, 50 mM Tris-HCl, pH 7) and subjected to lysis through incubation at 95 °C for 10 minutes. The lysates were then stored at -80 °C. For the polyphosphate extraction, 250 μL of 95% ethanol was added to each GITC-lysed sample, followed by vigorous mixing. This mixture was applied to a silica membrane spin column and centrifuged for 30 seconds at 16,100 x g. The flow-through was discarded, and 750 μL of a solution containing 5 mM Tris-HCl (pH 7.5), 50 mM NaCl, 5 mM EDTA, and 50% ethanol was added before centrifuging for 30 seconds at 16,100 x g. Once again, the flow-through was discarded, and a 2-minute centrifugation at 16,100 x g was carried out. The column was placed in a clean 1.5 mL microfuge tube, and 150 μL of 50 mM Tris-HCl (pH 8) was added. Following a 5-minute incubation at room temperature, polyP was eluted by centrifuging for 2 minutes at 8,000 x g. PolyP was quantified using the Phosfinity-Quant kit from Aminoverse, employing the scPPX enzyme.

### 7. Determination of the minimal inhibitory concentration (MIC) for colistin by broth microdilution assays

The colistin resistance of SGH10 WT, *Δppk1, Δppx, Δppk1-Δppx* was tested at concentrations ranging from 0.125 to 32 ug/mL. To do this, liquid cultures were grown in cation-adjusted Müller-Hinton broth (MH-2) from isolated colonies and incubated overnight at 37°C until reaching the stationary phase. Subsequently, 96-well plates were filled with 200 μL of MH-2 broth, supplemented with various concentrations of this antibiotic through serial dilutions in a 2-fold manner. Bacteria were then inoculated into the wells. Finally, the cultures were incubated for 32 hours with agitation at 37°C. During this period, bacterial growth was determined by measuring optical density at 600 nm using a Tecan M200 Infinite Pro microplate reader. MIC was defined as the lowest antibiotic concentration with no bacterial growth. *Pseudomonas aeruginosa* spp. PAO1 strain was used as a control for MIC determination.

### 8. Capsule production quantification

Capsule production was quantified by measuring the total amount of uronic acids, following the method previously described for *K. pneumoniae* ^5^. Briefly, uronic acids were extracted from 500 mL of culture using zwittergent, precipitated with ethanol, and resuspended in a tetraborate/sulfuric acid solution. Then, 3-phenyl phenol was added, and the uronic acid content was estimated by measuring the absorbance at 520 nm and comparing it to a standard curve generated with commercial glucuronic acid.

### 9. Low-speed centrifugation of bacteria

Four milliliters of a 10^9^ CFU/mL solution of the desired strain in LB were prepared in 15 mL Falcon tubes. The tubes were centrifuged at 2,000 × g for 10 min and imaged against a black background. The supernatant’s OD600nm was measured using a BioMateTM spectrophotometer (Thermo Scientific).

### 10. Quantification of Biofilm Biomass and EPS and Visualization Using Confocal Microscopy

Biofilm formation was assessed using a previously described protocol ^92^, with some modifications. Briefly, *K. pneumoniae* SGH10 wild-type and polyP mutant strains were grown overnight in LB broth, then diluted 1:100 in fresh LB broth. Next, 100 μL of each dilution was added to a 96-well polystyrene microtiter plate and incubated at 37°C for 24 h. Wells containing only media were used as blanks. After removing planktonic cells, wells were washed twice with sterile water and stained with 150 μL of 0.1% crystal violet for 30 min. Then, wells were rinsed twice with sterile water, and the stained biofilms were solubilized with 95% ethanol. Biofilm formation was quantified by measuring the OD595nm using a TECAN infinite 200 pro plate reader. To quantify EPS components such as curli and cellulose, the Ebbabiolight 680 probe was used because it bounds to these components without influencing biofilm formation. EbbaBiolight 680 was diluted in a growth medium at a 1:1000 ratio for the biofilm assay. Then, the supplemented growth medium was inoculated with the bacterial culture. Next, the wells of a 96-well plate were filled with 100 μL of the inoculated medium. The unused wells were filled with sterile water to prevent drying during incubation. The plate was covered with a lid or an adhesive seal. Fluorescence was measured at 540ex/680em in a TECAN infinite 200 pro plate reader. Each sample was measured in triplicate, and we calculated averages of the absorbance values for analysis. To investigate the accumulation of polyP in the biofilm and its influence on formation, an overnight culture was diluted 1:100 in the LB broth medium. Subsequently, a slide was submerged in 10 mL of the 1:100 culture inside a 50-mL falcon tube and incubated at 37°C for 24 h. The slide was washed three times with nanopure water and incubated for 30 minutes with 5 µg/mL DAPI to stain DNA and PolyP. The polyP accumulation was visualized using a CLSM Zeiss 757 confocal microscope, with a 358ex/461em laser to observe DNA and a 358ex/540em laser to observe polyP.

### 11. Scanning electron microscopy

To examine potential modifications at the superficial level, including the capsule and overall morphology, we conducted scanning electron microscopy (SEM) ^34^ with some changes. We prepared a lawn on LB agar from strains frozen at -80°C, then incubated for 24 h at 37°C. Using a sterile swab, we collected enough bacteria and resuspended them in PBS 1X. We adjusted the OD600nm to 2.8 (10^9^ CFU/ml) and fixed 500 µL of the suspension with 2.5% glutaraldehyde and 0.15% ruthenium red to improve the capsule visualization contrast. Samples were subjected to critical point drying and gold coating and subsequently observed in a high vacuum Zeiss EVO M15 scanning electron microscope. To examine superficial morphology in the mutants and WT strains following nutrient deprivation, overnight cultures of *K. pneumoniae* SGH10 and polyP mutants were diluted into fresh LB broth and cultured at 180 rpm and 37°C for 2 h. The bacterial cells were centrifuged twice in MOPS low-phosphate minimal medium (containing 0.4% glucose and 0.1 mM potassium phosphate) and incubated at 37°C for 2 h to stimulate polyP accumulation. The samples were treated in the same manner before observation using SEM.

### 12. Predation resistance assay

In this predation resistance assay, we semiquantitatively evaluated the possible attenuation of virulence in polyP metabolism mutants compared to the SGH10 WT strain. We modified a previously established protocol ^41^. Bacterial colonies were taken from a -80°C stock to prepare overnight cultures, and 300 µL were then seeded onto a plate using a Digralsky loop to generate a bacterial lawn. The plate was left to dry and incubated for 24 h at 23°C. Meanwhile, *D. discoideum* was cultured in HL5 medium, and serial dilutions were prepared to obtain the following cell concentrations: 500,000 - 50,000 - 5,000 – 500 – 50 and 5 cells per 5 µL. The bacterial lawns were then spotted with 5 µL of the serial *D. discoideum* dilutions. The plates were allowed to dry and were incubated at 21°C for 6 days. Plaque formation was visually examined on days 3 and 6. Isolates that did not enable amoebae growth were considered virulent for the amoeba. Bacterial strains that exhibited social development with 500 *D. discoideum* cells or less were considered sensitive to predation ^42^.

### 13. Social Development Assays

For social development assays, we followed a published protocol ^17^. Overnight cultures of each bacterial strain (30 μL) were homogeneously included per well of a 24-well plate containing N agar (1 g peptone, 1 g glucose, 20 g agar in 1 L of 17 mM Soerensen phosphate buffer) and grown overnight at 23^◦^C. A drop of a cellular suspension corresponding to 10^4^ *D. discoideum* cells in HL5 was spotted in the middle of each well and the plates were further incubated at 23^◦^C for 6 days. The social development of amoebae was monitored daily for 6 days, and the phase reached was scored, being classified as “aggregation,” “elevation,” and “culmination.” A score of “1” was assigned when amoebae aggregated, forming a phagocytosis plate, “2” when all the well surface elevated structures such as worms or fingers were observed, and “3” when fruiting bodies were formed across all the well surfaces. Transitions among these three phases were scored, with half of the value corresponding to the closest next stage. The number of fruiting bodies on a 1 cm^2^ surface was also quantified for each strain under study and was compared with the commensal KpgE strain. Additionally, images of social development were captured on days 1, 2, and 3 using a Zeiss Stemi 305 stereomicroscope with a total magnification of 10X, coupled to a 5 MP OPTIKA camera.

### 14. Phagocytosis assays

A lawn was prepared on LB agar from strains GFP frozen at -80°C and incubated for 24 h at 37°C. Bacteria were collected using a sterile swab and resuspended in Sorensen 1X buffer. Cell density was adjusted to OD600nm at 10^7^ cells/mL. Axenic cultures of *D. discoideum* were set to a concentration of 10^6^ cells/mL and 1 mL of culture was deposited in 24-well plates. The plate was centrifuged at 600 x g for 10 minutes and incubated at 23°C for 24 h. Each well was washed 3 times with Sorensen buffer and 1 mL of bacterial suspension was added to each well to obtain an MOI of 10. The plate was centrifuged at 600 x g for 30 min. The plate was then observed with the Lionheart FX automatic microscope, programmed to take photographs every 10 minutes for 24 to 48 h in bright field (BF) and GFP and in 4 regions of interest per well at 23°C. The images were analyzed with Gen5 software version 3.08 (Lionheart FX) and ImageJ software version 1.52 (Fiji version).

## AUTHOR INFORMATION

### Corresponding Author

*, Tel.:+56229787185, Laboratorio de Microbiología de Sistemas, Las Palmeras 3425, Ñuñoa CP 7800003, Santiago, Chile.

### Author Contributions

FC and AM conceived the study. DR, FC, AM, MV, MG, JV, and MD designed the experiments. DR and FC analyzed the data and interpreted the results. YG and YC developed the methodology for mutant generation. DR and MD carried out the mutant generation. DR, MD, and JV conducted the experiments. MH, CV, GN, EK, and PS performed proteomic processing and analysis. DR and FC wrote the manuscript. FC, AM, NG, and MH critically reviewed the manuscript. All the authors approved the final version of the manuscript.

### Founding Sources

This work was funded by the FONDECYT projects F. Chavez 1211852, and A. Marcoleta 1221193. Additionally, it received contributions from the FONDEQUIP project Lionheart EQM180216, and support from the María Ghilardi Foundation to D. Rojas.

## ACKNOWLEDGMENT

We express our gratitude to Nicole Molina for her support in preparing the required materials, and to Dr. T. Shiba for generously providing us with polyP of various sizes: short, medium, and large.

## Notes

The authors declare no competing financial interest.

## REFERENCES

1. Choby JE, Howard-Anderson J, Weiss DS. Hypervirulent *Klebsiella pneumoniae* – clinical and molecular perspectives. J Intern Med. 2020;287(3):283–300. doi:10.1111/joim.13007

2. Paczosa MK, Mecsas J. *Klebsiella pneumoniae*: Going on the Offense with a Strong Defense. Microbiology and Molecular Biology Reviews. 2016;80(3):629–661. doi:10.1128/mmbr.00078-15

3. Gonzalez-Ferrer S, Peñaloza HF, Budnick JA, et al. Finding Order in the Chaos: Outstanding Questions in *Klebsiella pneumoniae* Pathogenesis. Infect Immun. 2021;89(4). doi:10.1128/IAI.00693-20

4. Wyres KL, Holt KE. *Klebsiella pneumoniae* as a key trafficker of drug resistance genes from environmental to clinically important bacteria. Curr Opin Microbiol. 2018;45:131–139. doi:10.1016/j.mib.2018.04.004

5. Walker KA, Treat LP, Sepúlveda VE, Miller VL. The Small Protein RmpD Drives Hypermucoviscosity in *Klebsiella pneumoniae*. Heran Darwin K, ed. mBio. 2020;11(5). doi:10.1128/mBio.01750-20

6. Ovchinnikova OG, Treat LP, Teelucksingh T, et al. Hypermucoviscosity Regulator RmpD Interacts with Wzc and Controls Capsular Polysaccharide Chain Length. mBio. Published online May 4, 2023. doi:10.1128/mbio.00800-23

7. Russo TA, Olson R, MacDonald U, Beanan J, Davidsona BA. Aerobactin, but not yersiniabactin, salmochelin, or enterobactin, enables the growth/survival of hypervirulent (hypermucoviscous) *Klebsiella pneumoniae* ex vivo and in vivo. Infect Immun. 2015;83(8):3325–3333. doi:10.1128/IAI.00430-15

8. Struve C, Bojer M, Krogfelt KA. Characterization of *Klebsiella pneumoniae* type 1 fimbriae by detection of phase variation during colonization and infection and impact on virulence. Infect Immun. 2008;76(9):4055–4065. doi:10.1128/IAI.00494-08

9. Wang H, Wilksch JJ, Chen L, Tan JWH, Strugnell RA, Gee ML. Influence of Fimbriae on Bacterial Adhesion and Viscoelasticity and Correlations of the Two Properties with Biofilm Formation. Langmuir. 2017;33(1):100–106. doi:10.1021/acs.langmuir.6b03764

10. Wu MC, Lin TL, Hsieh PF, Yang HC, Wang JT. Isolation of Genes Involved in Biofilm Formation of a *Klebsiella pneumoniae* Strain Causing Pyogenic Liver Abscess. PLoS One. 2011;6(8):e23500. doi:10.1371/journal.pone.0023500

11. Zheng J xin, Lin Z wei, Chen C, et al. Biofilm Formation in *Klebsiella pneumoniae* Bacteremia Strains Was Found to be Associated with CC23 and the Presence of wcaG. Front Cell Infect Microbiol. 2018;8. doi:10.3389/fcimb.2018.00021

12. Russo TA, Marr CM. Hypervirulent *Klebsiella pneumoniae*. Clin Microbiol Rev. 2019;32(3). doi:10.1128/CMR.00001-19

13. Xu Q, Yang X, Chan EWC, Chen S. The hypermucoviscosity of hypervirulent *K. pneumoniae* confers the ability to evade neutrophil-mediated phagocytosis. Virulence. 2021;12(1):2050–2059. doi:10.1080/21505594.2021.1960101

14. Wyres KL, Lam MMC, Holt KE. Population genomics of *Klebsiella pneumoniae*. Nat Rev Microbiol. 2020;18(6):344–359. doi:10.1038/s41579-019-0315-1

15. Lam MMC, Wyres KL, Duchêne S, et al. Population genomics of hypervirulent *Klebsiella pneumoniae* clonal-group 23 reveals early emergence and rapid global dissemination. Nat Commun. 2018;9(1):2703. doi:10.1038/s41467-018-05114-7

16. Kornberg A, Rao NN, Ault-Riché D. Inorganic polyphosphate: a molecule of many functions. Annu Rev Biochem. 1999;68(1):89–125. doi:10.1146/annurev.biochem.68.1.89

17. Varas MA, Riquelme-Barrios S, Valenzuela C, et al. Inorganic polyphosphate is essential for *Salmonella Typhimurium* Virulence and survival in *Dictyostelium discoideum*. Front Cell Infect Microbiol. 2018;8(JAN). doi:10.3389/fcimb.2018.00008

18. Xie L, Jakob U. Inorganic polyphosphate, a multifunctional polyanionic protein scaffold. Journal of Biological Chemistry. 2019;294(6):2180–2190. doi:10.1074/jbc.REV118.002808

19. Rashid MH, Kornberg A. Inorganic polyphosphate is needed for swimming, swarming, and twitching motilities of Pseudomonas aeruginosa. Proc Natl Acad Sci U S A. 2000;97(9):4885–4890.

20. Fraley CD, Rashid MH, Lee SSK, et al. A polyphosphate kinase 1 (PPK1) mutant of *Pseudomonas aeruginosa* exhibits multiple ultrastructural and functional defects. Proc Natl Acad Sci U S A. 2007;104(9):3526–3531.

21. Ortiz-Severín J, Varas M, Bravo-Toncio C, Guiliani N, Chávez FP. Multiple antibiotic susceptibility of polyphosphate kinase mutants (PPK1 and PPK2) from *Pseudomonas aeruginosa* PAO1 as revealed by global phenotypic analysis. Biol Res. 2015;48:1–6. doi:10.1186/s40659-015-0012-0

22. Kim KS, Rao NN, Fraley CD, Kornberg A. Inorganic polyphosphate is essential for long-term survival and virulence factors in *Shigella* and *Salmonella* spp. Proceedings of the National Academy of Sciences. 2002;99(11):7675–7680. doi:10.1073/pnas.112210499

23. Varas MA, Riquelme-Barrios S, Valenzuela C, et al. Inorganic polyphosphate is essential for *Salmonella Typhimurium* Virulence and survival in Dictyostelium discoideum. Front Cell Infect Microbiol. 2018;8(JAN):1–18. doi:10.3389/fcimb.2018.00008

24. Rijal R, Cadena LA, Smith MR, Carr JF, Gomer RH. Polyphosphate is an extracellular signal that can facilitate bacterial survival in eukaryotic cells. Proceedings of the National Academy of Sciences. 2020;117(50):31923–31934. doi:10.1073/pnas.2012009117

25. Bowlin MQ, Gray MJ. Inorganic polyphosphate in host and microbe biology. Trends Microbiol. 2021;29(11):1013–1023. doi:10.1016/j.tim.2021.02.002

26. Roberge N, Neville N, Douchant K, et al. Broad-Spectrum Inhibitor of Bacterial Polyphosphate Homeostasis Attenuates Virulence Factors and Helps Reveal Novel Physiology of *Klebsiella pneumoniae* and *Acinetobacter baumannii*. Front Microbiol. 2021;12. doi:10.3389/fmicb.2021.764733

27. Chu WHW, Tan YH, Tan SY, et al. Acquisition of regulator on virulence plasmid of hypervirulent *Klebsiella* allows bacterial lifestyle switch in response to iron. mBio. Published online August 2, 2023. doi:10.1128/mbio.01297-23

28. Lv H, Zhou Y, Liu B, et al. Polyphosphate Kinase Is Required for the Processes of Virulence and Persistence in *Acinetobacter baumannii*. Microbiol Spectr. 2022;10(4). doi:10.1128/spectrum.01230-22

29. Aschar-Sobbi R, Abramov AY, Diao C, et al. High sensitivity, quantitative measurements of polyphosphate using a new DAPI-based approach. J Fluoresc. 2008;18(5):859–866. doi:10.1007/s10895-008-0315-4

30. Dahl JU, Gray MJ, Bazopoulou D, et al. The anti-inflammatory drug mesalamine targets bacterial polyphosphate accumulation. Nat Microbiol. 2017;2(January). doi:10.1038/nmicrobiol.2016.267

31. Zhu J, Wang T, Chen L, Du H. Virulence Factors in Hypervirulent *Klebsiella pneumoniae*. Front Microbiol. 2021;12(April):1–14. doi:10.3389/fmicb.2021.642484

32. Catalán-Nájera JC, Garza-Ramos U, Barrios-Camacho H. Hypervirulence and hypermucoviscosity: Two different but complementary Klebsiella spp. phenotypes Virulence. 2017;8(7):1111–1123. doi:10.1080/21505594.2017.1317412

33. Walker KA, Miner TA, Palacios M, et al. A *Klebsiella pneumoniae* regulatory mutant has reduced capsule expression but retains hypermucoviscosity. mBio. 2019;10(2):1–16. doi:10.1128/MBIO.00089-19

34. Tan YH, Chen Y, Chu WHW, Sham LT, Gan YH. Cell envelope defects of different capsule-null mutants in K1 hypervirulent *Klebsiella pneumoniae* can affect bacterial pathogenesis. Mol Microbiol. 2020;113(5):889–905. doi:10.1111/mmi.14447

35. Phanphak S, Georgiades P, Li R, King J, Roberts IS, Waigh TA. Super-Resolution Fluorescence Microscopy Study of the Production of K1 Capsules by *Escherichia coli*: Evidence for the Differential Distribution of the Capsule at the Poles and the Equator of the Cell. Langmuir. 2019;35(16):5635–5646. doi:10.1021/acs.langmuir.8b04122

36. Rendueles O. Deciphering the role of the capsule of *Klebsiella pneumoniae* during pathogenesis: A cautionary tale. Mol Microbiol. 2020;113(5):883–888. doi:10.1111/mmi.14474

37. Fraley CD, Rashid MH, Lee SSK, et al. A polyphosphate kinase 1 (*ppk1*) mutant of *Pseudomonas aeruginosa* exhibits multiple ultrastructural and functional defects. Proceedings of the National Academy of Sciences. 2007;104(9):3526–3531. doi:10.1073/pnas.0609733104

38. Rashid MH, Rumbaugh K, Passador L, et al. Polyphosphate kinase is essential for biofilm development, quorum sensing, and virulence of *Pseudomonas aeruginosa*. Proceedings of the National Academy of Sciences. 2000;97(17):9636–9641. doi:10.1073/pnas.170283397

39. Choong FX, Huzell S, Rosenberg M, et al. A semi high-throughput method for real-time monitoring of curli producing Salmonella biofilms on air-solid interfaces. Biofilm. 2021;3:100060. doi:10.1016/j.bioflm.2021.100060

40. Ho JY, Lin TL, Li CY, et al. Functions of Some Capsular Polysaccharide Biosynthetic Genes in *Klebsiella pneumoniae* NTUH K-2044. PLoS One. 2011;6(7):e21664. doi:10.1371/journal.pone.0021664

41. Filion G, Charette SJ. Assessing Pseudomonas aeruginosa virulence using a nonmammalian host: *Dictyostelium discoideum*. Methods in Molecular Biology. 2014;1149:671–680. doi:10.1007/978-1-4939-0473-0_51

42. Paquet VE, Charette SJ. Amoeba-resisting bacteria found in multilamellar bodies secreted by *Dictyostelium discoideum:* social amoebae can also package bacteria. FEMS Microbiol Ecol. 2016;92(3):fiw025. doi:10.1093/femsec/fiw025

43. Bravo-Toncio C, Álvarez JA, Campos F, et al. *Dictyostelium discoideum* as a surrogate host–microbe model for antivirulence screening in *Pseudomonas aeruginosa* PAO1. Int J Antimicrob Agents. Published online 2016:1–7. doi:10.1016/j.ijantimicag.2016.02.005

44. Marcoleta AE, Varas MA, Ortiz-Severín J, et al. Evaluating different virulence traits of *Klebsiella pneumoniae* using *Dictyostelium discoideum* and zebrafish larvae as host models. Front Cell Infect Microbiol. 2018;8(FEB). doi:10.3389/fcimb.2018.00030

45. Wang L, Shen D, Wu H, Ma Y. Resistance of hypervirulent *Klebsiella pneumoniae* to both intracellular and extracellular killing of neutrophils. PLoS One. 2017;12(3):e0173638. doi:10.1371/journal.pone.0173638

46. Rasko DA, Sperandio V. Anti-virulence strategies to combat bacteria-mediated disease. Nat Rev Drug Discov. 2010;9(2):117–128. doi:10.1038/nrd3013

47. Cegelski L, Marshall GR, Eldridge GR, Hultgren SJ. The biology and future prospects of antivirulence therapies. Nat Rev Microbiol. 2008;6(1):17–27.

48. DIckey SW, Cheung GYC, Otto M. Different drugs for bad bugs: Antivirulence strategies in the age of antibiotic resistance. Nat Rev Drug Discov. Published online 2017. doi:10.1038/nrd.2017.23

49. Downey M. A stringent analysis of polyphosphate dynamics in *Escherichia coli*. J Bacteriol. 2019;201(9):1–6. doi:10.1128/JB.00070-19

50. Gray MJ. Inorganic polyphosphate accumulation in *Escherichia coli* is regulated by DksA but not by (p)ppGpp. J Bacteriol. 2019;201(9):1–20. doi:10.1128/JB.00664-18

51. Bravo-Toncio C, Álvarez JA, Campos F, et al. *Dictyostelium discoideum* as a surrogate host–microbe model for antivirulence screening in *Pseudomonas aeruginosa* PAO1. Int J Antimicrob Agents. Published online 2016:1–7. doi:10.1016/j.ijantimicag.2016.02.005

52. Palacios M, Miner TA, Frederick DR, et al. Identification of two regulators of virulence that are conserved in *Klebsiella pneumoniae* classical and hypervirulent strains. mBio. 2018;9(4). doi:10.1128/mBio.01443-18

53. Tinsley CR, Manjula BN, Gotschlich EC. Purification and characterization of polyphosphate kinase from *Neisseria meningitidis*. Infect Immun. 1993;61(9):3703–3710. doi:10.1128/iai.61.9.3703-3710.1993

54. Müller WEG, Schröder HC, Wang X. Inorganic Polyphosphates As Storage for and Generator of Metabolic Energy in the Extracellular Matrix. Chem Rev. 2019;119(24):12337–12374. doi:10.1021/acs.chemrev.9b00460

55. Varas M, Valdivieso C, Mauriaca C, et al. Multi-level evaluation of *Escherichia coli* polyphosphate related mutants using global transcriptomic, proteomic and phenomic analyses. Biochimica et Biophysica Acta (BBA) - General Subjects. 2017;1861(4):871–883. doi:10.1016/j.bbagen.2017.01.007

56. Meredith TC, Woodard RW. Characterization of *Escherichia coli* <SCP>D</SCP> - arabinose 5-phosphate isomerase encoded by *kpsF*: implications for group 2 capsule biosynthesis. Biochemical Journal. 2006;395(2):427–432. doi:10.1042/BJ20051828

57. Perepelov A V., Ujazda E, Senchenkova SN, Shashkov AS, Kaca W, Knirel YA. Structural and serological studies on the O-antigen of Proteus mirabilis O14, a new polysaccharide containing 2-[(R)-1-carboxyethylamino]ethyl phosphate. Eur J Biochem. 1999;261(2):347–353. doi:10.1046/j.1432-1327.1999.00251.x

58. Element SJ, Moran RA, Beattie E, Hall RJ, van Schaik W, Buckner MMC. Growth in a biofilm promotes conjugation of a *bla* _NDM-1_-bearing plasmid between *Klebsiella pneumoniae* strains. mSphere. Published online July 7, 2023. doi:10.1128/msphere.00170-23

59. Vuotto C, Longo F, Balice MP, Donelli G, Varaldo PE. Antibiotic resistance related to biofilm formation in *Klebsiella pneumoniae*. Pathogens. 2014;3(3):743–758. doi:10.3390/pathogens3030743

60. Wang G, Zhao G, Chao X, Xie L, Wang H. The characteristic of virulence, biofilm and antibiotic resistance of *Klebsiella pneumoniae*. Int J Environ Res Public Health. 2020;17(17):1–17. doi:10.3390/ijerph17176278

61. Chhibber S, Gondil VS, Sharma S, Kumar M, Wangoo N, Sharma RK. A novel approach for combating *Klebsiella pneumoniae* biofilm using histidine functionalized silver nanoparticles. Front Microbiol. 2017;8(JUN):1–10. doi:10.3389/fmicb.2017.01104

62. Alcántar-Curiel MD, Ledezma-Escalante CA, Jarillo-Quijada MD, et al. Association of Antibiotic Resistance, Cell Adherence, and Biofilm Production with the Endemicity of Nosocomial *Klebsiella pneumoniae*. Biomed Res Int. 2018;2018. doi:10.1155/2018/7012958

63. Schumachera MA, Zenga W. Structures of the activator of *k. pneumonia* biofilm formation, mrkh, indicates pilz domains involved in c-di-gmp and DNA binding. Proc Natl Acad Sci U S A. 2016;113(36):10067–10072. doi:10.1073/pnas.1607503113

64. Dos Santos Goncalves M, Delattre C, Balestrino D, et al. Anti-biofilm activity: A function of *Klebsiella pneumoniae* capsular polysaccharide. PLoS One. 2014;9(6). doi:10.1371/journal.pone.0099995

65. Chen T, Dong G, Zhang S, et al. Effects of iron on the growth, biofilm formation and virulence of *Klebsiella pneumoniae* causing liver abscess. BMC Microbiol. 2020;20(1). doi:10.1186/s12866-020-01727-5

66. Shi X, Rao NN, Kornberg A. Inorganic polyphosphate in Bacillus cereus: Motility, biofilm formation, and sporulation. Proc Natl Acad Sci U S A. 2004;101(49):17061–17065. doi:10.1073/pnas.0407787101

67. Drozd M, Chandrashekhar K, Rajashekara G. Polyphosphate-mediated modulation of *Campylobacter jejuni* biofilm growth and stability. Virulence. 2014;5(6):680–690. doi:10.4161/viru.34348

68. Recalde A, van Wolferen M, Sivabalasarma S, Albers SV, Navarro CA, Jerez CA. The role of polyphosphate in motility, adhesion, and biofilm formation in sulfolobales. Microorganisms. 2021;9(1):1–13. doi:10.3390/microorganisms9010193

69. Rashid MH, Rumbaugh K, Passador L, et al. Polyphosphate kinase is essential for biofilm development, quorum sensing, and virulence of *Pseudomonas aeruginosa*. Proc Natl Acad Sci U S A. 2000;97(17):9636–9641. doi:10.1073/pnas.170283397

70. Grillo-Puertas M, Villegas JM, Rintoul MR, Rapisarda VA. Polyphosphate Degradation in Stationary Phase Triggers Biofilm Formation via LuxS Quorum Sensing System in *Escherichia coli*. PLoS One. 2012;7(11):e50368. doi:10.1371/journal.pone.0050368

71. Ballén V, Gabasa Y, Ratia C, Sánchez M, Soto S. Correlation Between Antimicrobial Resistance, Virulence Determinants and Biofilm Formation Ability Among Extraintestinal Pathogenic *Escherichia coli* Strains Isolated in Catalonia, Spain. Front Microbiol. 2022;12. doi:10.3389/fmicb.2021.803862

72. Luo C, Chen Y, Hu X, et al. Genetic and Functional Analysis of the *pks* Gene in Clinical *Klebsiella pneumoniae* Isolates. Microbiol Spectr. Published online June 21, 2023. doi:10.1128/spectrum.00174-23

73. Tang-Fichaux M, Chagneau C V., Bossuet-Greif N, Nougayrède JP, Oswald É, Branchu P. The Polyphosphate Kinase of Escherichia coli Is Required for Full Production of the Genotoxin Colibactin. mSphere. 2020;5(6). doi:10.1128/mSphere.01195-20

74. Beaufay F, Quarles E, Franz A, Katamanin O, Wholey WY, Jakob U. Polyphosphate Functions *In Vivo* as an Iron Chelator and Fenton Reaction Inhibitor. mBio. 2020;11(4). doi:10.1128/mBio.01017-20

75. Erskine E, MacPhee CE, Stanley-Wall NR. Functional Amyloid and Other Protein Fibers in the Biofilm Matrix. J Mol Biol. 2018;430(20):3642–3656. doi:10.1016/j.jmb.2018.07.026

76. Bieler S, Estrada L, Lagos R, Baeza M, Castilla J, Soto C. Amyloid Formation Modulates the Biological Activity of a Bacterial Protein. Journal of Biological Chemistry. 2005;280(29):26880–26885. doi:10.1074/jbc.M502031200

77. Marcoleta AE, Berríos-Pastén C, Nuñez G, Monasterio O, Lagos R. *Klebsiella pneumoniae* Asparagine tDNAs Are Integration Hotspots for Different Genomic Islands Encoding Microcin E492 Production Determinants and Other Putative Virulence Factors Present in Hypervirulent Strains. Front Microbiol. 2016;7. doi:10.3389/fmicb.2016.00849

78. Flemming HC, Wingender J, Szewzyk U, Steinberg P, Rice SA, Kjelleberg S. Biofilms: an emergent form of bacterial life. Nat Rev Microbiol. 2016;14(9):563–575. doi:10.1038/nrmicro.2016.94

79. Cano V, March C, Insua JL, et al. *Klebsiella pneumoniae* survives within macrophages by avoiding delivery to lysosomes. Cell Microbiol. 2015;17(11):1537–1560. doi:10.1111/cmi.12466

80. Froquet R, Lelong E, Marchetti A, Cosson P. *Dictyostelium discoideum*: a model host to measure bacterial virulence. Nat Protoc. 2009;4(1):25–30. doi:10.1038/nprot.2008.212

81. Fung CP, Chang FY, Lin JC, et al. Immune response and pathophysiological features of *Klebsiella pneumoniae* liver abscesses in an animal model. Laboratory Investigation. 2011;91(7):1029–1039. doi:10.1038/labinvest.2011.52

82. Joseph L, Merciecca T, Forestier C, Balestrino D, Miquel S. From *Klebsiella pneumoniae* Colonization to Dissemination: An Overview of Studies Implementing Murine Models. Microorganisms. 2021;9(6):1282. doi:10.3390/microorganisms9061282

83. Thayil SM, Morrison N, Schechter N, Rubin H, Karakousis PC. The role of the novel exopolyphosphatase MT0516 in *Mycobacterium tuberculosis* drug tolerance and persistence. PLoS One. 2011;6(11). doi:10.1371/journal.pone.0028076

84. Shi X, Rao NN, Kornberg A. Inorganic polyphosphate in *Bacillus cereus*: Motility, biofilm formation, and sporulation. Proc Natl Acad Sci U S A. 2004;101(49):17061–17065. http://www.pubmedcentral.nih.gov/articlerender.fcgi?artid=535361&tool=pmcentrez&rendertype=abstract

85. Tinsley CR, Manjula BN, Gotschlich EC. Purification and characterization of polyphosphate kinase from *Neisseria meningitidis*. Infect Immun. Published online 1993. doi:10.1128/iai.61.9.3703-3710.1993

86. Kreppel L. dictyBase: a new *Dictyostelium discoideum* genome database. Nucleic Acids Res. 2004;32(90001):332D–333. doi:10.1093/nar/gkh138

87. Basu S, Fey P, Pandit Y, Dodson R, Kibbe WA, Chisholm RL. dictyBase 2013: integrating multiple Dictyostelid species. Nucleic Acids Res. 2012;41(D1):D676–D683. doi:10.1093/nar/gks1064

88. Fey P, Dodson RJ, Basu S, Chisholm RL. One Stop Shop for Everything *Dictyostelium*: dictyBase and the Dicty Stock Center in 2012. In:; 2013:59–92. doi:10.1007/978-1-62703-302-2_4

89. Fey P, Kowal AS, Gaudet P, Pilcher KE, Chisholm RL. Protocols for growth and development of *Dictyostelium discoideum*. Nat Protoc. 2007;2(6):1307–1316. doi:10.1038/nprot.2007.178

90. Li H, Durbin R. Fast and accurate short read alignment with Burrows–Wheeler transform. Bioinformatics. 2009;25(14):1754–1760. doi:10.1093/bioinformatics/btp324

91. Pokhrel A, Lingo JC, Wolschendorf F, Gray MJ. Assaying for Inorganic Polyphosphate in Bacteria. Journal of Visualized Experiments. 2019;(143). doi:10.3791/58818

92. Chen T, Dong G, Zhang S, et al. Effects of iron on the growth, biofilm formation and virulence of *Klebsiella pneumoniae* causing liver abscess. BMC Microbiol. 2020;20(1):1–7. doi:10.1186/s12866-020-01727-5

