## Supplementary material for "Effects of Polyphosphate metabolism Mutations on Biofilm, Capsule Formation and Virulence Traits in Hypervirulent ST23 *Klebsiella pneumoniae* SGH10"

**This PDF file includes:** Supplementary figures S1-S8

### Supplementary Figures

**Supplementary table 1.** Strains, plasmids and primers used in this study

| Bacterial strain or plasmid | Genotype or comments | Source/Reference |
|---|---|---|
| *Klebsiella aerogenes* (KpgE) DBS0305928 | Recommended strain for supporting the growth of D. discoideum | Dicty Stock Center |
| *Escherichia coli* S17 lpir | Laboratory strain typically used for cloning purpose | Laboratory Collection |
| *Klebsiella pneumoniae* SGH10 | This strain was isolated from a patient's hepatic abscess in 2014 and was found to be hypermucoviscous and hypervirulent. | Andres Marcoleta`s laboratory collection |
| *K. pneumoniae* SGH10 D*ppk1* | Null mutant for the gene that encodes the enzyme that synthesizes inorganic polyphosphate. | This work |
| *K. pneumoniae* SGH10 D*ppx* | Null mutant for the gene that encodes the enzyme that degrades inorganic polyphosphate | This work |
| *K. pneumoniae* SGH10 D*ppk*D*ppx* | Double null mutant for the operon that metabolizes polyphosphate (polyP). | This work |
| *K. pneumoniae* SGH10 D*wcaJ* | Null mutant for the gene encoding the WcaJ glycosyltransferase that performs the first step in capsule polysaccharide synthesis. Strain used as a control. | Andres Marcoleta`s laboratory collection |
| pBBR1-GFP | Constitutive expression of the GFP protein, $Kan^r$ | |
| pR6KtetsacB | | |

| | |
|---|---|
| PPK1_UP_F | TGACATGATTACGAATTCTGCATGAAAAACATCGTGGT |
| PPK1_UP_R | GGGGTGTTATCGTTTAACCGCTCACACTCCGTTTTA |
| PPK1_DOWN_F | ACGGAGTGTGAGCGGTTAAACGATAACACCCCTCGGC |
| PPK1_DOWN_R | AAGCTTGCATGCCTGCAGGCGGCTACGAATATCCTGGT |
| PPX_UP_F | ACATGATTACGAATTCAGGATTCGCGCATTATTGAC |
| PPX_UP_R | GTTGATTGCCAGAGATTGTTCGAACCAAGGTCAACC |
| PPX_down_F | GACCTTGGTTCGAACAATCTCTGGCAATCAACCGAG |
| ppx_down_R | AAGCTTGCATGCCTGCAGGCCGGTCAAGCATCTGTATT |
| CTR_PPK1_Fw | 5´-CCGACGAATGAACAACCAG-3´ |
| CTR_PPK1_Rv | 5´-CAACGCCCATAAAAATCAGG-3` |
| CTR_PPX_Fw | 5`-TCAGTATTGTGGACCGTTTCC-3` |
| CTR_PPX_Rv | 5`-CTACGCCCGGTTAAAAACAA-3 |
| CTR_PoliP_Fw | 5´-CCGACGAATGAACAACCAG-3´ |
| CTR_PoliP_Rv | 5`-CTACGCCCGGTTAAAAACAA-3 |

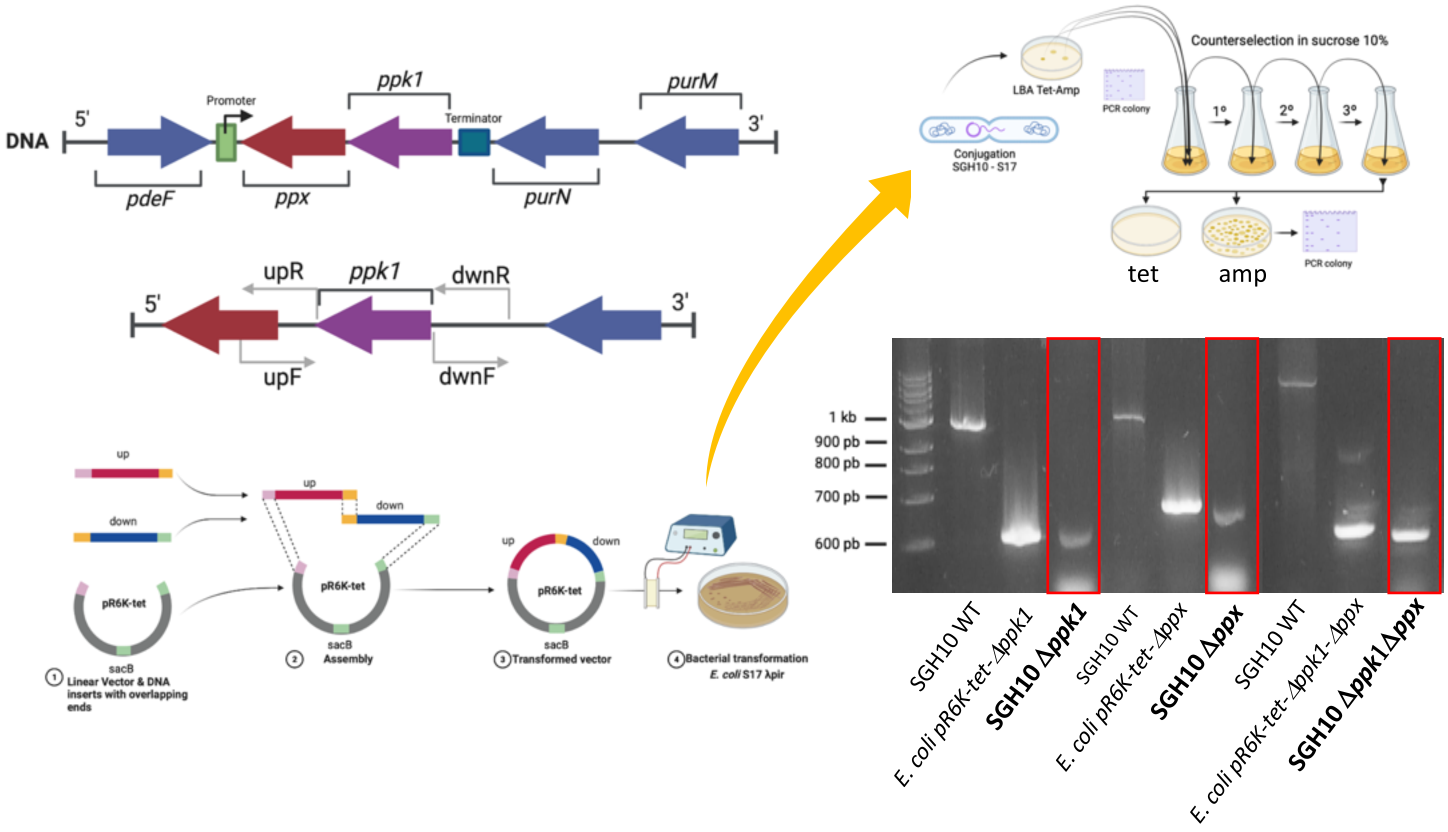


**Supplementary Fig 1. Workflow in mutant development.** First, we amplified ~1000 bp fragments upstream and downstream of the target gene from genomic DNA templates using Q5 high-fidelity DNA polymerase (New England Biolabs) and assembled them into the vector pR6KTet-SacB using NEBuilder® HiFi DNA Assembly Master Mix (New England Biolabs). The resulting plasmids were then introduced into *K. pneumoniae* SGH10 through conjugation with the *E. coli* donor strain S17-1λpir. After selecting single cross-transconjugants in medium containing Tetracycline (50 µg/ml), we eliminated the donor E. coli by adding ampicillin (100 µg/ml) to the medium. To counteract selection of the sacB gene in the pR6KTet-sacB backbone, single tetracycline-resistant crosses were passaged in LB medium without sodium chloride but supplemented with 20% sucrose. Successful deletion of the gene of interest was confirmed by selecting tetracycline-sensitive double crossovers via polymerase chain reaction. We also confirmed the mutant strains, including Δ*ppk1*, Δ*ppx*, and Δ*ppk1*-Δ*ppx*, through scar generation in the SGH10 genome by sequencing the fragments and matching them with the complete genome of the SGH10 WT.

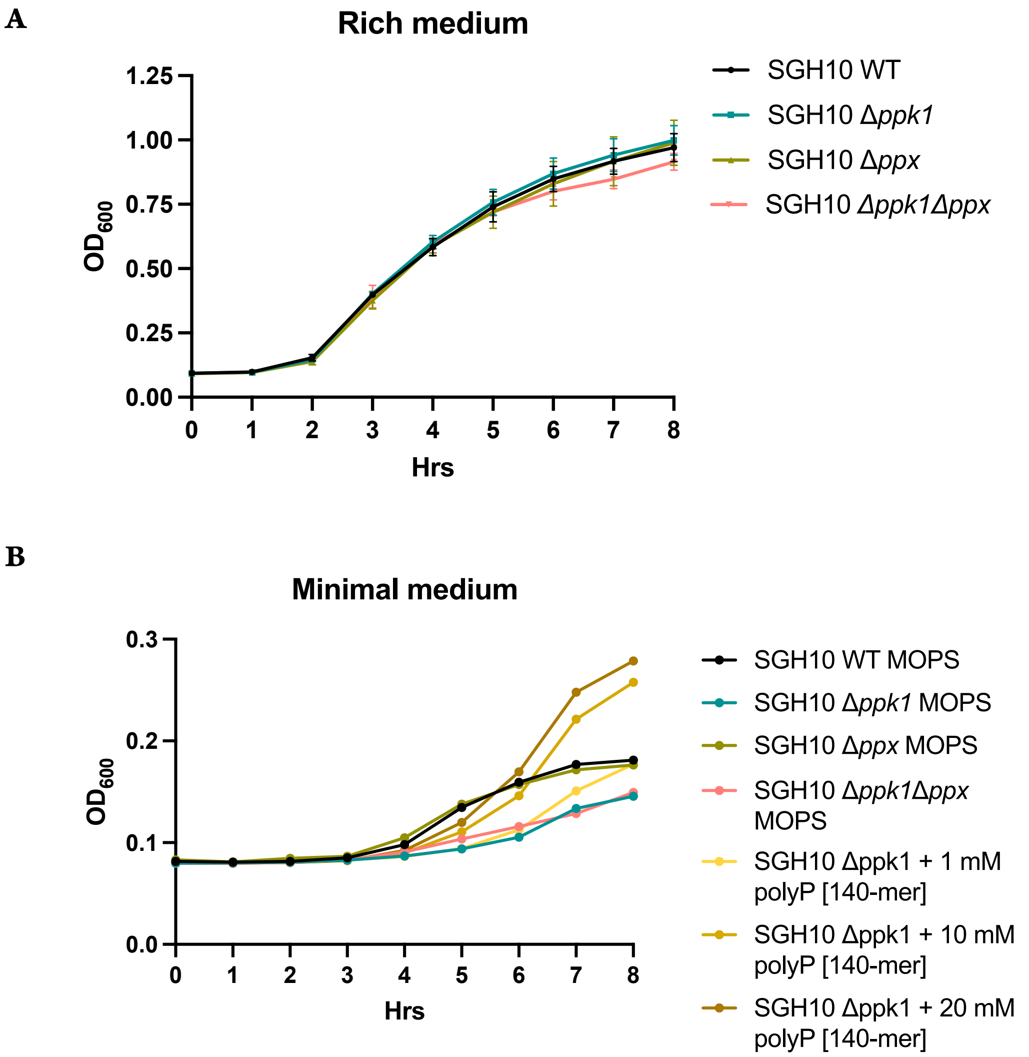


**Supplementary Fig 2. Growth curve.** Mutant strains in polyP metabolism and SGH10 WT were grown overnight. A 1:1000 dilution in LB **(A)** or **(B)** MOPS was inoculated in a 96-well plate (MOPS culture medium was inoculated with 4 g/l glucose and 0.1 mM $H_2PO_4^-$). 1, 10 and 20 mM of polyP [140-mer] were used for phenotype reconstitution

It was incubated for 24 hrs in a TECAN infinite 200 pro plate reader at 37ºC and 180 rpm. Readings were taken every 10 minutes (n=6) to $DO_{600}$

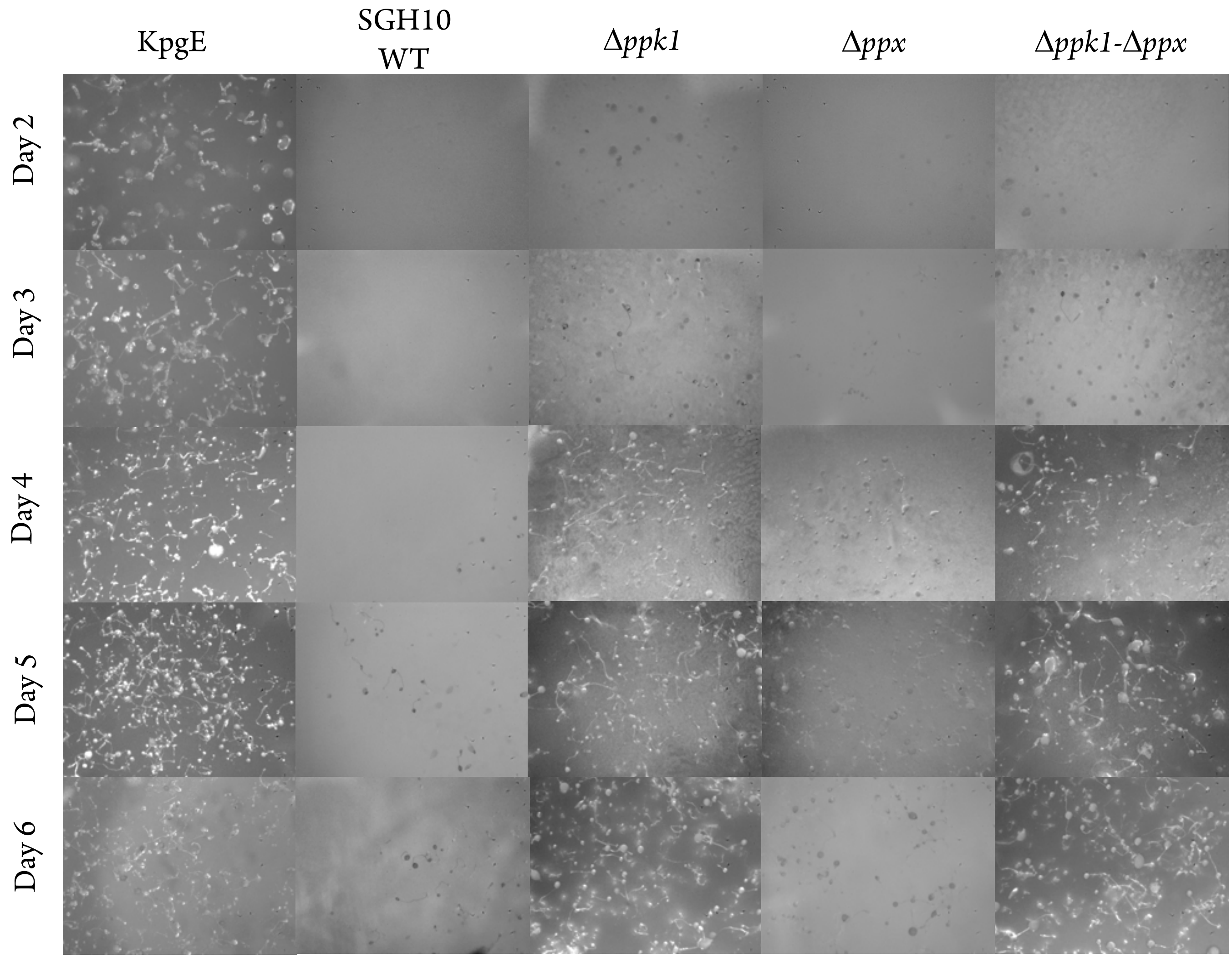


**Supplementary Fig 3. Social development assay.** 30 uL of overnight cultures of the strains under study were inoculated in a 24-well plate on N agar to generate a lawn. They were incubated for 24 hrs at 23 ºC. *D. discoideum* was adjusted to $1x10^6$/ml. A 10 uL drop of the dilution was inoculated on each lawn. The social development of the amoeba was followed for six days. The KpgE commensal strain was used as a control.

**A**

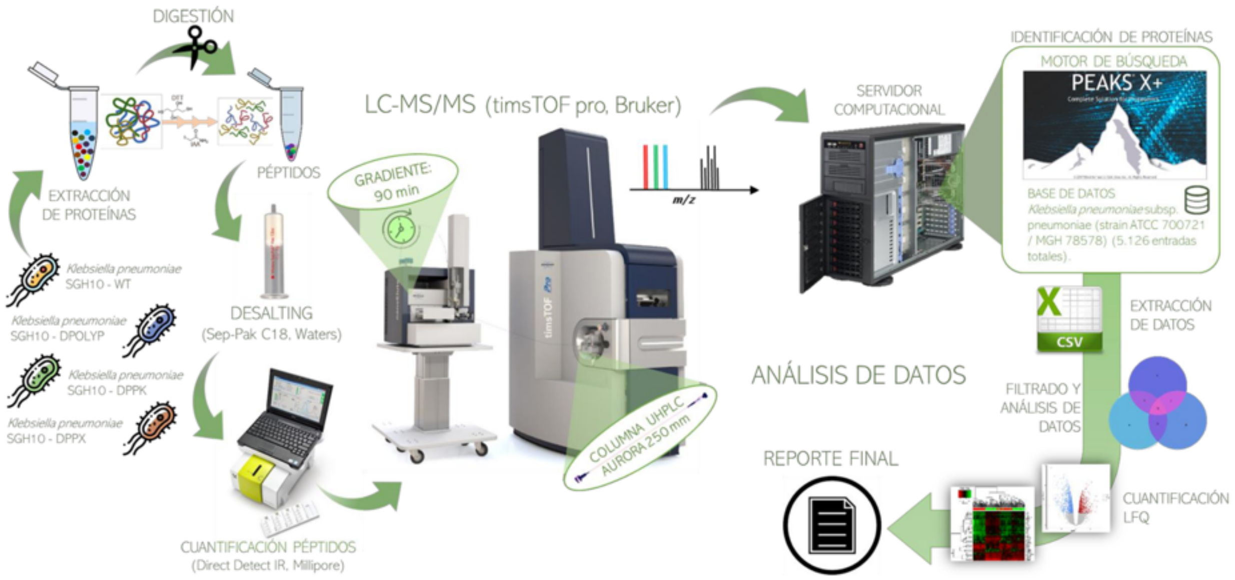


**B**

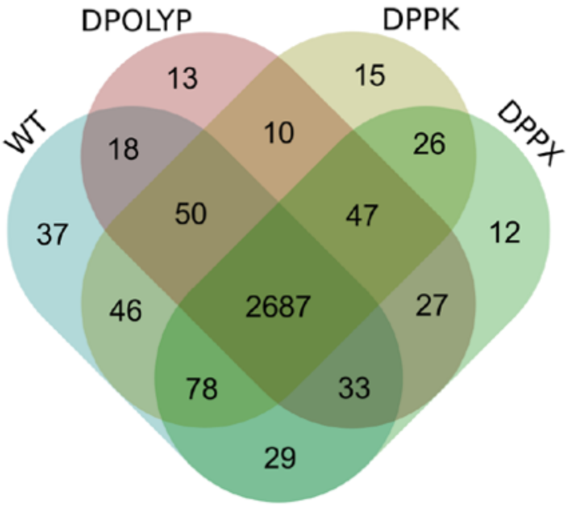


**C**

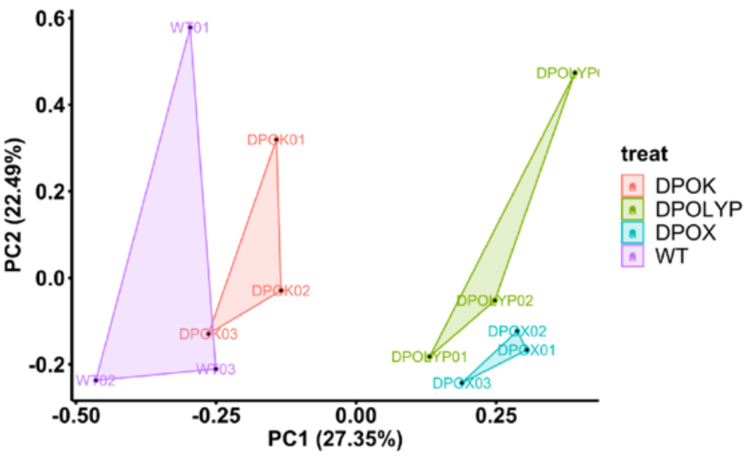


**Supplementary Fig 4.** **(A)** Diagram of the workflow used in protein identification. **(B)** Comparison of the protein group from the sum of all proteins per biological replicate depending on each treatment received. **(C)** Principal Component Analysis (PCA) of the set of proteins identified per sample.

| **Comparison** | **Proteins** | |
|---|---|---|
| | Quantifiable | DEPs (FDR < 0,05) |
| D*ppk1* vs WT | 2.287 | 302 |
| D*ppx* vs WT | | 728 |
| D*ppk1*D*ppx* vs WT | | 673 |

**Supplementary Table 2.** Number of quantifiable proteins and DEPs (Differential Expression Proteins) obtained for each of the comparisons.

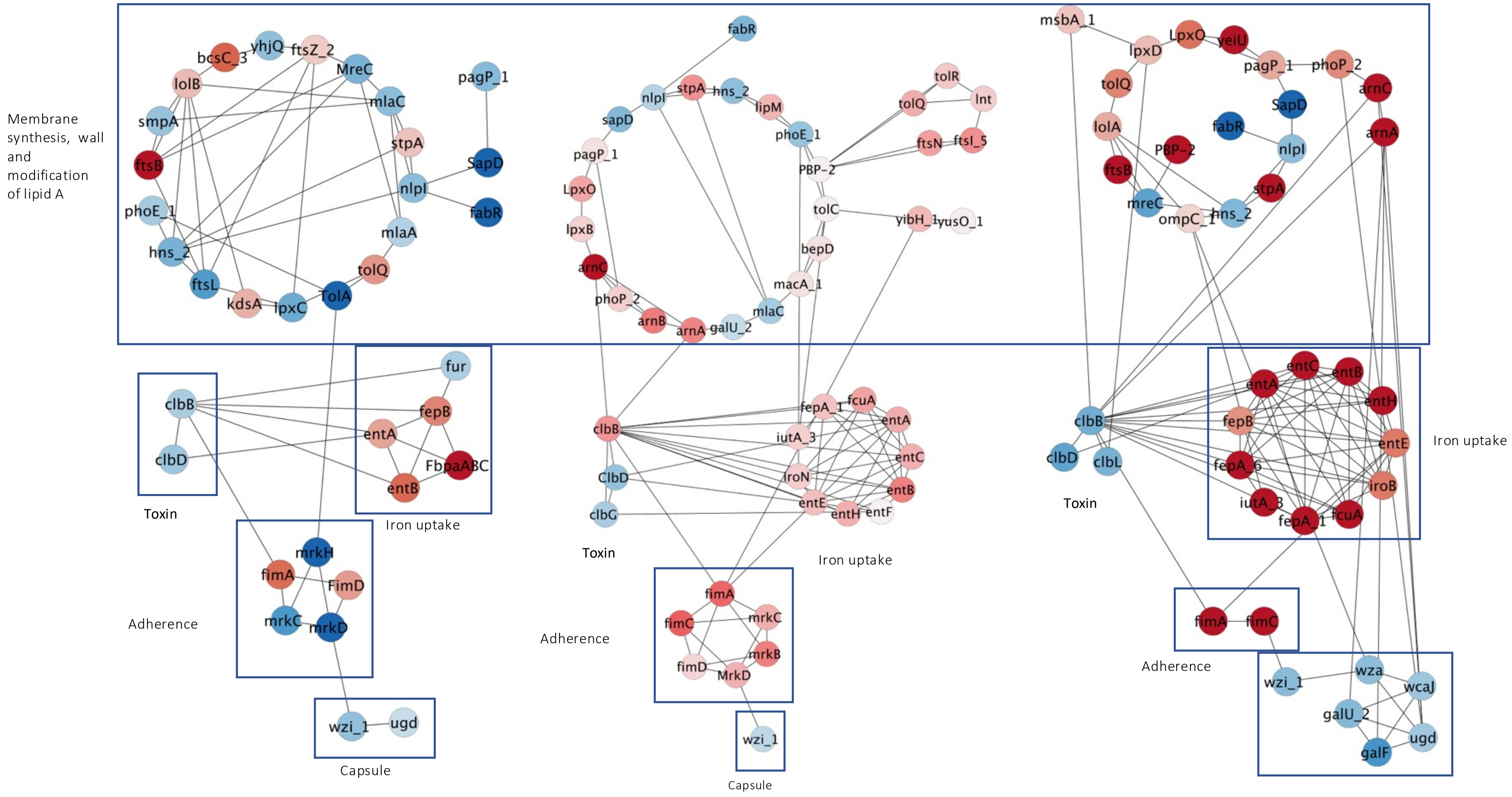

Membrane synthesis, wall and modification of lipid A
Toxin
Iron uptake
Adherence
Capsule
Toxin
Iron uptake
Adherence
Capsule
Toxin
Iron uptake
Adherence

**Supplementary Fig 5.** Network of curated interactions between proteins involved in the virulence of Klebsiella pneumoniae. Clusters of proteins related to capsule formation and biofilms are observed; siderophore synthesis; formation of membrane, wall and modification of lipid A; antibiotic resistance. The network was generated in Cytoscape.

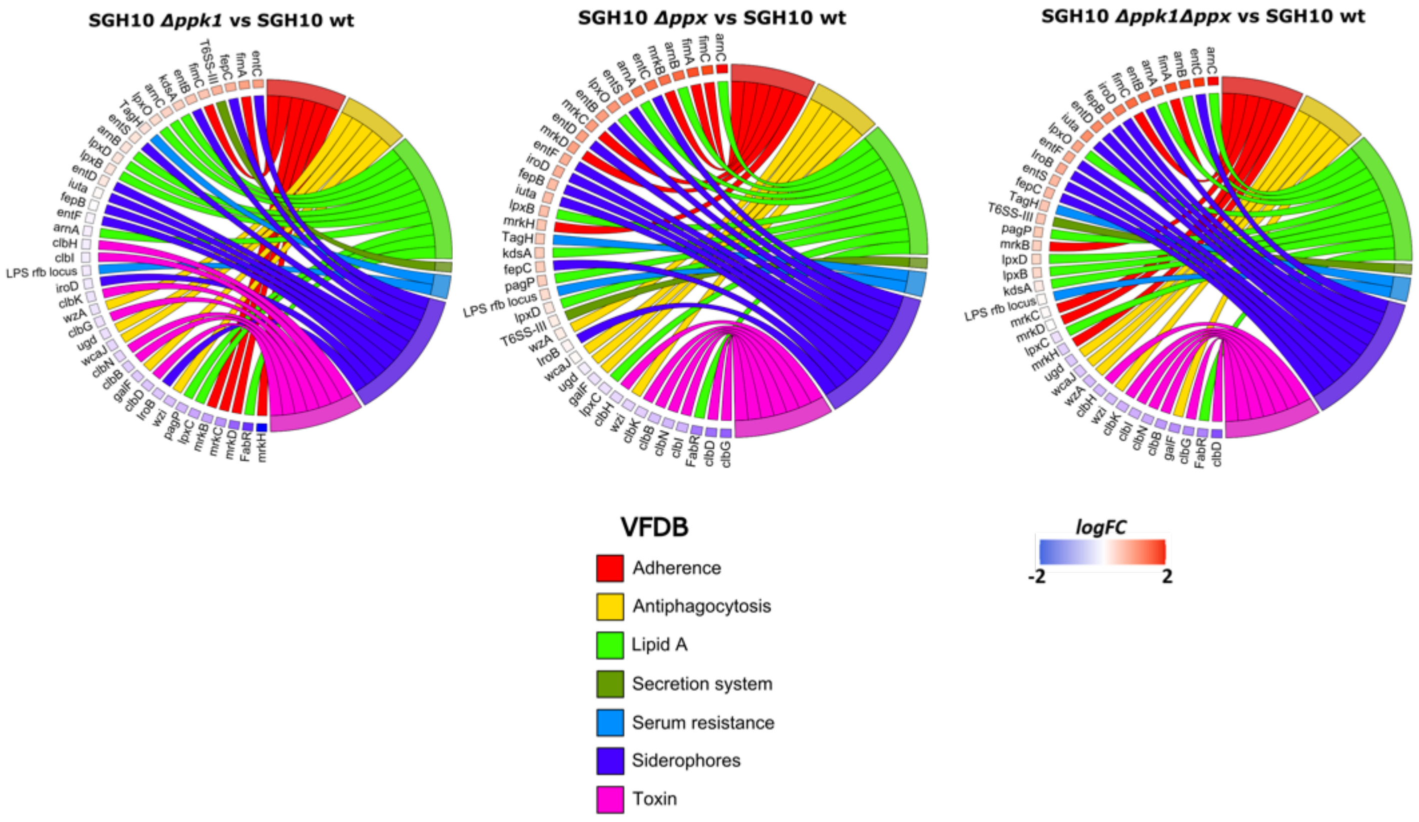


**Supplementary Figure 6:** Interaction Chordplot

The chordplot graph illustrates protein-protein interactions categorized based on the VFDB database. Each arc corresponds to an interaction category, while the external nodes represent the identified proteins. The color of each node corresponds to the LogFC value from each analysis.

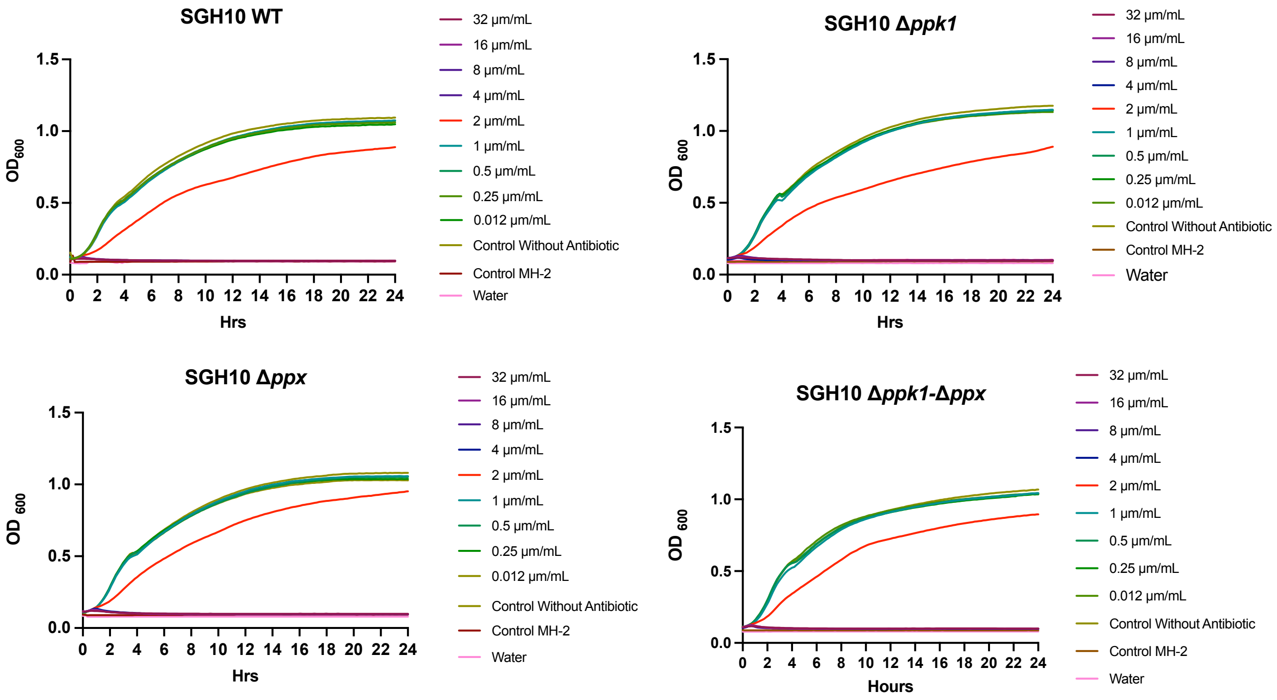


**Supplementary Figure 7:** Colistin resistance assessment of SGH10 strains, Δ*ppk1,* Δ*ppx,* Δ*ppk1*-Δ*ppx,* and others at concentrations ranging from 0.125 to 32 ug/mL using liquid cultures. The MIC (minimum inhibitory concentration), the lowest antibiotic concentration inhibiting bacterial growth, was determined with Pseudomonas aeruginosa spp. PAO1 as a control.
